# Geometry Can Provide Long-Range Mechanical Guidance for Embryogenesis

**DOI:** 10.1101/075309

**Authors:** Mahamar Dicko, Pierre Saramito, Guy B. Blanchard, Claire M. Lye, Bénédicte Sanson, Jocelyn Étienne

## Abstract

Downstream of gene expression, effectors such as the actomyosin contractile machinery drive embryo morphogenesis. During *Drosophila* embryonic axis extension, actomyosin has a specific planar-polarised organisation, which is responsible for oriented cell intercalation. In addition to these cell rearrangements, cell shape changes also contribute to tissue deformation. While cell-autonomous dynamics are well described, understanding the tissue-scale behaviour challenges us to solve the corresponding mechanical problem at the scale of the whole embryo, since mechanical resistance of all neighbouring epithelia will feedback on individual cells. Here we propose a novel numerical approach to compute the whole-embryo dynamics of the actomyosin-rich apical epithelial surface. We input in the model specific patterns of actomyosin contractility, such as the planar-polarisation of actomyosin in defined ventro-lateral regions of the embryo. Tissue strain rates and displacements are then predicted over the whole embryo surface according to the global balance of stresses and the material behaviour of the epithelium. Epithelia are modelled using a rheological law that relates the rate of deformation to the local stresses and actomyosin anisotropic contractility. Predicted flow patterns are consistent with the cell flows observed when imaging *Drosophila* axis extension *in toto*, using light sheet microscopy. The agreement between model and experimental data indicates that the anisotropic contractility of planar-polarised actomyosin in the ventro-lateral germband tissue can directly cause the tissue-scale deformations of the whole embryo. The three-dimensional mechanical balance is dependent on the geometry of the embryo, whose curved surface is taken into account in the simulations. Importantly, we find that to reproduce experimental flows, the model requires the presence of the cephalic furrow, a fold located anteriorly of the extending tissues. The presence of this geometric feature, through the global mechanical balance, guides the flow and orients extension towards the posterior end.

**Author Summary:** The morphogenesis of living organisms is a facinating process during which a genetic programme controls a sequence of molecular changes which will cause the original embryo to acquire a new shape. While we have a growing knowledge of the timing and spatial distribution of key molecules downstream of genetic programmes, there remain gaps of understanding on how these patterns can generate the appropriate mechanical force, so as to deform the tissues in the correct manner. In this paper, we show how a model of tissue mechanics can link the known pattern of actomyosin distribution in *Drosophila* tissues to the process of axis extension, which is a ubiquitous morphogenetic movement of developing animal embryos. We show in numerical simulations that the correct movement is obtained only if the geometry of the embryo presents some precise features. This means that prior morphogenetic movements responsible for these features need to have succeeded in order to carry on the next round of morphogenesis. This highlights the contribution of mechanical feedback on the morphogenetic programme and also how mechanical action integrates at the scale of the whole embryo.

## Introduction

The morphogenesis of living organisms involves precise shape changes and displacements of the tissues that constitute the embryo under the control of gene expression [1]. These movements result from changes in the mechanical balance, which can be caused by local growth [2] or by local activation of the contractile machinery of actomyosin [3]. The action of these effectors can integrate at the scale of a whole tissue through the establishment of a new mechanical balance, which happens quasi-instantaneously in the absence of inertia [4]. This integration at a tissue-scale explains how a local process can effect a global deformation [5–9]. The mechanical coupling of different tissues and extra-cellular structures crucially changes the resulting mechanical balance globally, and in turn can lead to very different morphologies. For instance, uncoupling the *Drosophila* pupal wing blade from the cuticle at its margin prevents its elongation during hinge contraction [8].

Axis extension of the *Drosophila* embryo at gastrulation offers a good system to model convergence and extension morphogenetic flows, which are ubiquitous in the development of animals [10]. During axis extension, a region of the epithelial monolayer that makes up the embryo, the ventro-lateral part of the germ-band, narrows in the dorsoventral (DV) direction and simultaneously extends in the orthogonal direction (antero–posterior, AP) towards the posterior end, Fig 1*a–c*. This movement is known to depend on the genetically specified organisation of actomyosin in a planar-polarised manner, with an enhanced Myosin II recruitment specifically along the cell–cell junctions aligned with the DV axis [11,12], Fig 2*a–c*. This planar-polarised actomyosin is responsible for active shortening of these DV junctions, which resolve in active cell intercalations [13]. The total tissue deformation of *Drosophila* axis extension is however not explained by the sole action of planar-polarised myosin [14]. In addition, the extending germ-band is subjected to an external pull originating from the invagination of the posterior midgut [6,7], Fig 1*a–c*. As a result, cells change shape, elongating along the AP axis, and the deformation of the tissue is the combination of both cell intercalation and cell shape strain contributions [14,15].

**Fig 1:**
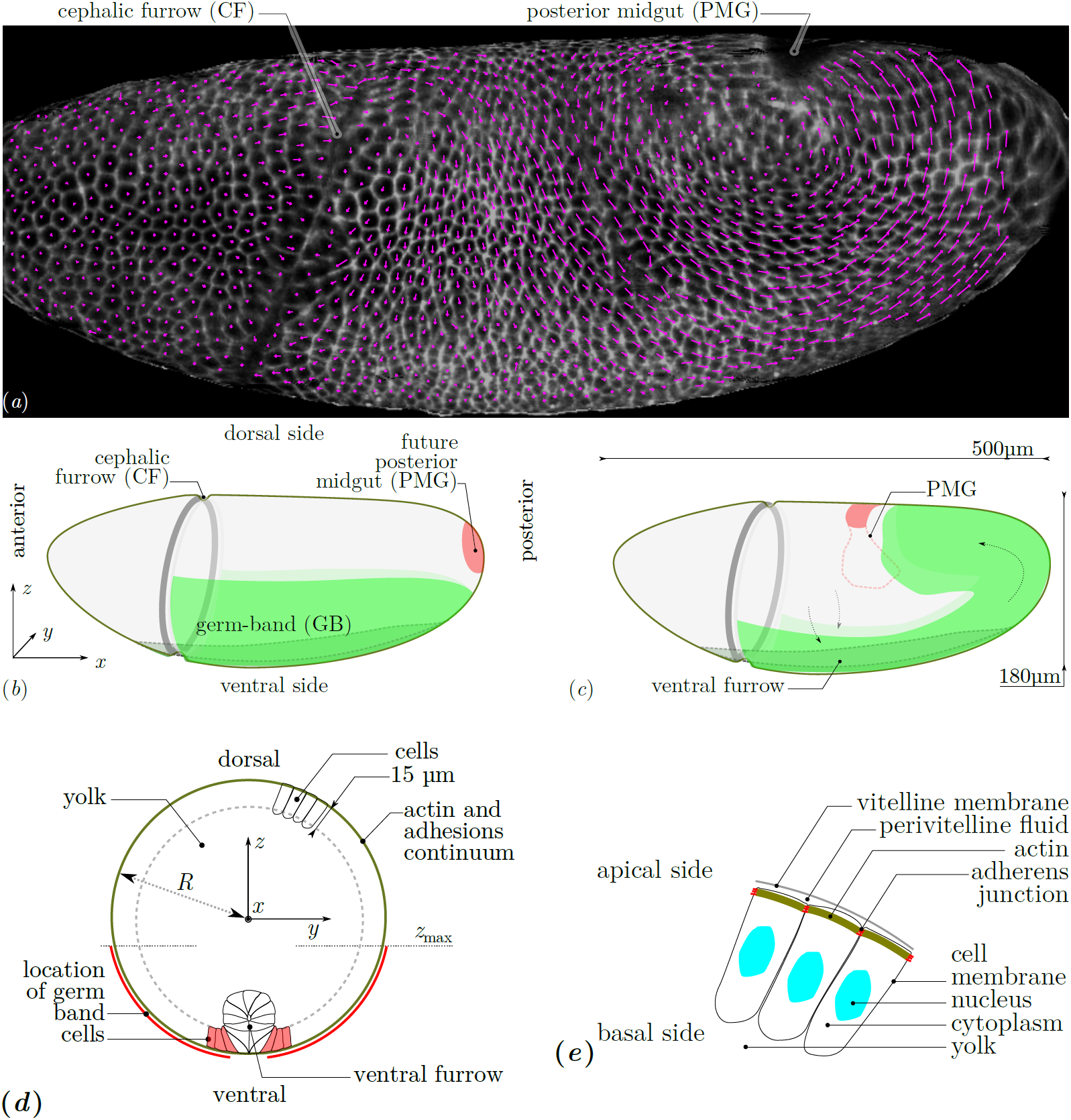
(*a*) Lateral view of a wildtype *Drosophila* embryo and flow of cells during GB extension. *White*, two-dimensional rendering of signal of membrane markers in an apical curved image layer of a three-dimensional *z*-stack acquired by light-sheet microscopy (specifically, mSPIM) at an arbitrary time instant (late fast phase of GB extension). *Magenta*, two-dimensional projection of the displacement of the centroid of each cell over the following 30 seconds. (*b*) Geometry and tissue configuration of *Drosophila* embryo [5] immediately prior to GB extension. Tissues situated at the outer surface are in solid colors, dashed lines correspond to structures internal to the embryo. (*c*) Sketch of morphogenetic movements and tissue configuration during GB extension and PMG invagination. (*d*) Geometry and structures of mechanical relevance in a transverse cut. The coordinate origin is in the centre. Contiguous cells form a continuous surface at the periphery of the embryo, the external limit is the cell’s apical side, the internal one (dashed line) their basal side. Only some cells are drawn. On the ventral side, a ventral furrow forms before GB extension and seals at the ventral midline just as the GB starts extending. Within the cells, actin structures form apically and are connected from one cell to the other by adhesive molecules, forming an embryo-scale continuum at the periphery of the embryo. The GB is highlighted in red, in this region myosin is activated in a planar-polarised manner. (*e*) Sketch of structures of mechanical relevance in the epithelial cells. The *vitelline membrane* is a rigid impermeable membrane. The *perivitelline fluid* is incompressible and viscous. The *actomyosin* of *Drosophila* cells is located at their apical surface, it is a thin layer (*<* 1 *µ*m) connected to other cell’s actomyosin via *adherens junctions*. The *cytoplasm* of cells behaves as an incompressible viscous fluid during the flow [32]. It is enclosed in the cell membranes, which have a low permeability but present excess area compared to cell’s volume. Beyond the basal surface of the cell monolayer, the *yolk* is an incompressible viscous fluid.

In this system, the dynamics of cell intercalation and myosin planar-polarisation have been modelled using discrete element approaches, which explicitly account for each cell–cell junction [16–19]. However, so far, it has not been possible in these modelling approaches to avoid using arbitrary boundary conditions to represent the mechanical resistance of the surrounding epithelia. The several morphogenetic movements occurring at once during *Drosophila* gastrulation interact through the global mechanical balance, and thus germ-band extension and posterior midgut invagination are not independent of one another [6,7]. In addition to the germ-band and posterior midgut, the neighbouring dorsal tissue, the future amnioserosa, is also being deformed during axis extension, although there is no active process reported in this tissue [5,9]. In this paper, we choose to focus on the tissue scale dynamics and the importance of the global mechanical balance on morphogenetic flows. To understand this, we solve a mechanical problem set on a three-dimensional surface that has the shape of the embryo. We treat the apical surfaces of the epithelial cells as a single continuum. Indeed, in the course of a convergence–extension process for an isolated tissue, the dynamics governing the tissue-scale can be captured by a material law involving only the resistance to shear and the energetic cost of area variations, see Fig 2*e–h*. These energetic costs sum per unit area the average cost of the deformations of each cell and the one of their rearrangement, which are the two components of tissue strain [15]. This coarse-grained view implies that the results of the simulation do not distinguish in what measure a given tissue deformation is achieved through cell intercalation or through cell shape change, Fig 2*f–g*.

**Fig 2:**
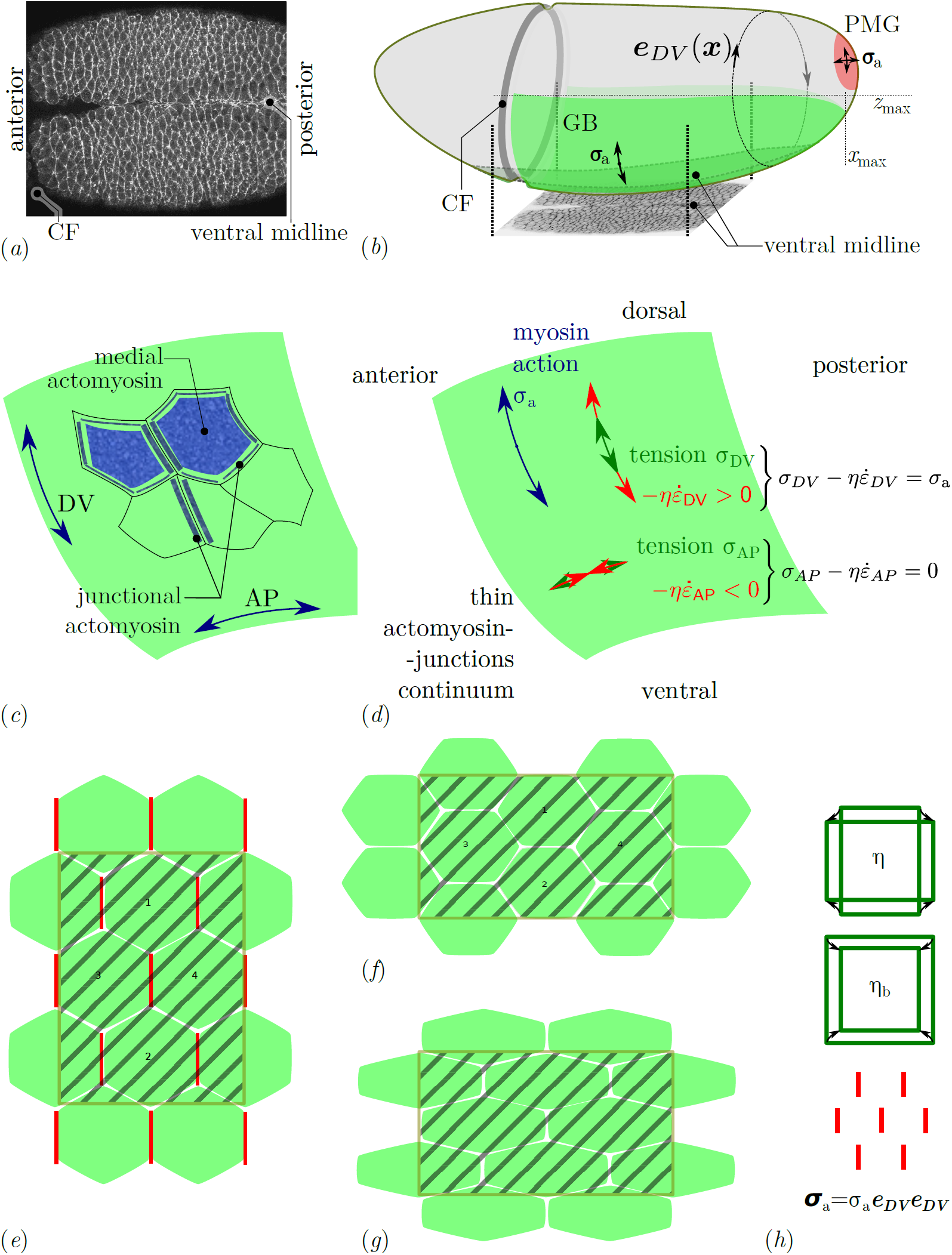
Myosin distribution during GB extension. (*a*) Fluorescently labeled myosin in the GB and midline over a ventral region early in GB extension [5]. The myosin is significantly denser along DV-oriented cell junctions (*y* direction) than along AP-oriented ones (*x* direction). This planar polarisation can be quantified [11,19,41]. (*b*) Sketch of the geometry of the entire embryo with the planar-polarised GB region (*green*) and the isotropically contracting PMG region (*red*) [5]. The region shown in panel *a* is shown from below. Isotropic contraction is assumed to be linked with an isotropic action of myosin, thus ***σ*****_a_** is an isotropic tensor in the PMG, whereas planar polarisation results in an anisotropic prestress ***σ*****_a_** [21], whose orientation we take as ***e***_DV_ *⊗ **e***_DV_, where ***e***_DV_ is a tangential unit vector orthogonal to the main axis of the embryo. (*c*) Sketch of the different pools of myosin present at the cell apices. Junctional myosin is associated with cell-cell junctions, and may form supracellular cables. Medial myosin is apical myosin not associated with junctions. (*d*) Tangential apical stresses in an arbitrary region of the GB. According to the constitutive relation, Eq (2), the (opposite of) viscous stress 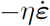 and mechanical stress ***σ*** need to balance the myosin prestress ***σ*****_a_** in both AP and DV directions. Since myosin prestress is zero along AP, the mechanical stress is equal to the viscous stress in this direction, 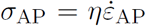, thus AP tension results in extension. In the DV direction, we have 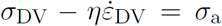, resulting in a combination of DV tension and contraction (convergence). The global mechanical balance, Eq (1), has to be solved in order to calculate ***σ*** and 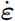. (*e*)–(*h*) Tissue strain, cell intercalation and cell shape change schematics (*e*) Initial cell arrangement, with planar-polarised myosin along DV-oriented cell-cell junctions (vertical) and the definition of a region of interest to track tissue deformation (*hashed area*) (*f*) A combination of cell intercalation (cells marked 1 and 2 are now neighbours) and cell shape and area changes leads to a tissue-scale deformation [15]. The region of interest has now a reversed aspect ratio along AP and DV, and has changed area. (*g*) A different combination of these cell-scale events (here no intercalation but more extensive pure shear of single cells and same area change) can lead to the same tissue-scale deformation. (*h*) Tissue scale resistance to deformation can be quantified by two numbers, a shear viscosity *η* and a second viscosity *η*_b_ corresponding to the additional resistance to area variations. Planar-polarised myosin prestress can be retained at tissue scale as an anisotropic prestress tensor ***σ*****_a_**.

At a continuum level, the mechanical effect of myosin contracting a meshwork of actin with a specific orientation can be modelled as a prestress with a specific orientation [20,21]. This modelling of anisotropic actomyosin stress generation has already been succesfully applied to tissues undergoing morphogenesis [22,23]. This allows us to ask the question whether the mechanism of intercalation is in itself the necessary step to convert the planar-polarised actomyosin activity into convergence and extension. Alternatively, the prestress generated by planar-polarised actomyosin may in itself cause tissue convergence and extension at a global scale, while local balances would govern how much of intercalation and how much of cell shape changes are incurred. In order to predict tissue-scale deformations, a mechanical model needs to have access to the reaction forces of any neighbouring tissue. The reaction forces of the other epithelia neighbouring the extending GB will depend on their own deformation. Therefore, it is necessary to include all of these epithelia in a mechanical approach. Here we propose a numerical technique that solves the flow generated by any patterning of actomyosin along the apical surface of the embryonic epithelia. Our modelling is based on some mechanical assumptions on how the apical actomyosin in the developing epithelium interacts with its environment, and on a rheological model of actomyosin itself [21]. The numerical simulation of flows on a curved surface is based on a novel finite element technique, which is not limited to potential flows as in previous literature [24], and for which we have proven accuracy properties [25].

Experimentally, it is now possible to image live embryos labelled with fluorescent proteins in toto, using light sheet imaging [26,27]. In *Drosophila*, this allowed experimentalists to observe morphogenetic events happening in different regions of the embryo and investigate how these impact on each other [6,9]. A current challenge is to quantify the corresponding deformations for the whole embryo as it has been done for smaller regions of tissue with limited curvature [6,14,15,28,29]. Another challenge is to develop numerical tools that permit the implementation of models over the whole embryo. Numerical methods taking into account the three-dimensional geometry of *Drosophila* embryo have been used to study the formation of the ventral furrow at gastrulation (reviewed in [30]) and germ-band extension [31]. However, the driving forces in the three-dimensional approaches were the observed cell-autonomous phenomenology rather than the patterning of actomyosin activation.

Here, without explicitly accounting for cell intercalations, we show that anisotropic myosin prestress can cause the global movements observed in *Drosophila* embryonic axis extension. We show that either a planar-polarisation of actomyosin in the germ-band or the pulling force due to the posterior midgut invagination are sufficient to generate a posterior-ward convergence and extension flow of the tissue, consistent with experimental evidence. Using existing movies of whole *Drosophila* embryos [6], we quantify cell flows and show that the numerical predictions are consistent with these. Finally, we show that a geometric feature of the embryo, the cephalic furrow, modifies the predicted flow and acts as a barrier for tissue deformation. This guides the convergence and extension flow towards the posterior of the embryo, therefore breaking the flow symmetry.

## Modelling

At the developmental stage of GB extension, the *Drosophila* embryo is made up of a single epithelial cell sheet that has an ovoid shape, with the cell apices facing outwards, see Fig 1. Gastrulation occurs immediately before GB extension, forming the ventral furrow, which begins to seal just at GB extension onset [6,14]. For simplicity, we will consider in this paper only the times that follow the completion of mesoderm sealing, thus cells can be considered to be mechanically connected across the ventral midline, Fig 1*d*. The apical actomyosin cortex is mechanically coupled from one cell to the other by transmembrane adherens junctions [33], which, away from specific folds of the epithelium, are located within 1 *µ*m of the apical surface [34].
From a mechanical point of view, the apical surface of the embryo can thus be seen as a thin layer of apical actomyosin seamed together by adherens junctions. Within this apical domain, the dynamics of active cell rearrangement and shape changes [13] and the causal planar-polarised recruitment of actomyosin [11,12] mediate GB extension. In consequence, our mechanical approach is based on this thin active layer which we model as a thin shell, similarly to what has been done for the cortex of single cells, in combination either with an elastic material law [35] or a viscoelastic one [21]. For a given active force generation, the dynamics of deformation of this apical meshwork is determined by its material properties and by the global mechanical balance [4], including interactions with neighbouring fluids and structures further apically and basally. On the basal side of this meshwork, baso-lateral cell membranes, cell cytoplasms and nuclei (Fig 1*d,e*) are not known to actively deform, and have been shown to flow as a viscous medium during gastrulation [32], immediately prior to GB extension. This passive behaviour implies that they are felt only as a drag (viscous friction). Note that there is no report of the presence of extra-cellular matrix on the basal side of this epithelium. On the outer side of the apical meshwork is the perivitelline membrane, but no specific adhesions bind them together, and the perivitelline liquid can play the role of a lubricating fluid between the two. In a first approximation, these two effects result in a friction force per unit area equal to the product of the velocity ***v*** by a friction coefficient *c*_f_. In terms of forces, the balance of the forces tangentially to the surface is thus:

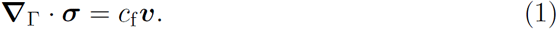
EQN1

The stress tensor ***σ*** is the tension in the apical shell of geometry Γ, and ***∇***_Γ_*·* is the surface divergence operator [36].

In order to obtain a *closed* model (i.e., a self-sufficient one), we need to supplement mechanical balance with a material law which links the stress to the deformations of the apical shell, and, in the present case, to the myosin activity also. We have recently derived and validated such a material law [21] by quantitative comparison of predictions of forces exerted by the actomyosin cortex of single cells with experimental measurements. The main ingredients of this modelling are the elastic resistance of actomyosin to deformation, which gives its short-term response, the turnover of actin and actin-crosslinking proteins ranging from seconds to minutes [37], which leads to a long-term liquid-like behaviour, and the myosin activity, which endows the cytoskeleton with emergent material properties [21]. It is worthwhile to note that, while the linear theory of transiently crosslinked gels does not lead to anisotropic material properties [38], the myosin activity generates an anisotropic stress, see supplementary material in [21]. For a tissue, the material ingredients are similar, with the addition of cell rearrangements which provide an alternative cause of elastic stress relaxation. We have shown that indeed, the same material law can describe successfully the rheology of early epithelia [39], although the value of parameters can vary. In a linear approximation, the apical actomyosin of embryos can thus be expected to have a viscoelastic liquid behaviour which can be written in the general form:

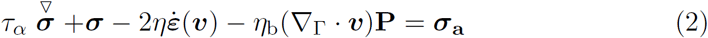
EQN2
where *τ_α_* is the relaxation time of actomyosin, 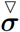 the objective derivative of the stress tensor, *η* and *η*_b_ are effective shear and compression viscosities of actomyosin,

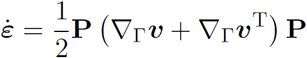
UEQN1
is the rate-of-strain tensor, and **P** is the projection tensor onto Γ, which is also the identity tensor on the surface. The tensor ***σ*****_a_** describes the myosin contractility, and can be understood as a prestress: because of myosin action, the meshwork of actin is continuously being offset from its stress-free configuration. The pre-stress is proportional to the myosin concentration and rate of power strokes [21,39]. The relaxation time *τ_α_* was found to be around 15 minutes in our previous work [21], which means that for the 90-minute germ band extension process, we are interested in times longer than relaxation. We have also shown for similar equations in another biophysical context and in one dimension [40] that the term *τ_α_* 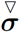 did not introduce marked qualitative features to the flow for time scales as short as the relaxation time itself. For the sake of simplicity, we will thus neglect the term *τ_α_* 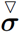 in what follows. Fig 2*h* illustrates the terms that sum to the tissue stress.

With this hypothesis, the mechanical balance equation (1) and the constitutive equation (2) can be combined into a single equation:

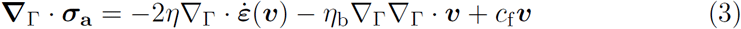
EQN3

On the left-hand side of Eq (3) is the myosin term, which provides the energy for the motion. On the right-hand side are two terms corresponding to energy dissipation: forces of friction with the actomyosin’s environment, and the viscous forces, corresponding to the cost of deforming the apical shell. These do not distinguish between inter-cellular dissipation, i.e. the mechanical energy spent during cell rearrangements in breaking cell-cell adhesive bonds and the one gained when establishing new ones; and intra-cellular dissipation, such as the cost of deforming the actomyosin cortex. Rather, these are lumped together, and taking a linear approximation, represented by effective viscosities *η* and *η*_b_.

The myosin contractility term ***σ*****_a_** is the source term that provides energy to the system and causes the deformations. We have shown theoretically and verified experimentally [21,39] that to a first approximation it is proportional to the local myosin concentration. However, when actomyosin recruitment is anisotropic, the tensor ***σ*****_a_** is also anisotropic, see supplementary materials in [21], and can be decomposed into a spatially-dependent intensity *σ*_a_ and an orientation tensor **A**, ***σ*****_a_** = *σ*_a_**A**. In the case of GB extension, it is observed experimentally [11,19,42] that myosin is activated in the GB, thus *σ*_a_ *>* 0 in the GB, and is recruited along DV-oriented cell junctions, thus **A** = ***e***_DV_ *⊗ **e***_DV_, see Fig 2.

Note that we impose mathematically that velocities are tangential with the embryo surface (see S1 Text). Indeed, this is what is observed experimentally, we have thus made the simplifying choice of assuming this rather than try to predict it. A model bypassing this hypothesis would need to be significantly more complicated, as the force balance in the normal direction would need to be calculated in addition to the one in the tangent plane. This force balance should include the pressure difference between the periviteline liquid and the embryo interior beyond the apical membranes (yolk and cytoplasms) and the viscous drag from these structures, and also the tension and bending forces in the cell apices. Although we do not model this, the force needed to keep the flow tangential in our simulations is calculated, and is the Lagrange multiplier associated with the tangentiality constraint.

In order to solve Eq (3), we use a finite-element approach. In [24], a related problem is addressed, and a finite-element method is described to solve surface-incompressible flow problems. This restriction to surface incompressibility allows the authors to reduce the problem to a PDE in terms of a scalar unknown, namely the stream function. Here we want a more generic approach allowing to address flows of finite surface-compressibility *η*_b_. We have thus developed a Lagrange-multiplier approach which can approximate the solution of vectorial equations set on a curved surface of 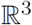 [25], see S1 Text. The resulting problem is discretised with finite elements using a triangle mesh of the embryo shape. The finite element free software rheolef is used to implement the method and its accuracy is verified on a test problem.

## Results

### A continuum shell model of the apical actomyosin cytoskeleton and adherens junctions meshwork can produce convergence and extension flows

Using the above model and the numerical technique described in S1 Text, we obtain numerical approximations of the solution of this problem for a geometry Γ closely mimicking the shape of *Drosophila* embryo. Fig 3 shows an example of a simulation result. The green colour codes *σ*_a_(***x***), the location where myosin is assumed to be activated along dorsoventral direction **A** = ***e**_DV_ ⊗ **e**_DV_*, and the arrows are the predicted velocity of the surface displacement of the apical continuum. This predicted flow is strongly dominated by two laterally-located vortices, which have their centre slightly dorsal from the edge of the GB region. They are rotating such that the velocity in the GB is strongly towards the posterior. The vortices are situated in the tissues located dorsally of the germband (future amnioserosa), which will thus be extensively deformed by the flow. This is indeed the case in *Drosophila* development, although part of the deformations *in vivo* may be related to PMG invagination [6], which is not included in this simulation (see below).

**Fig 3:**
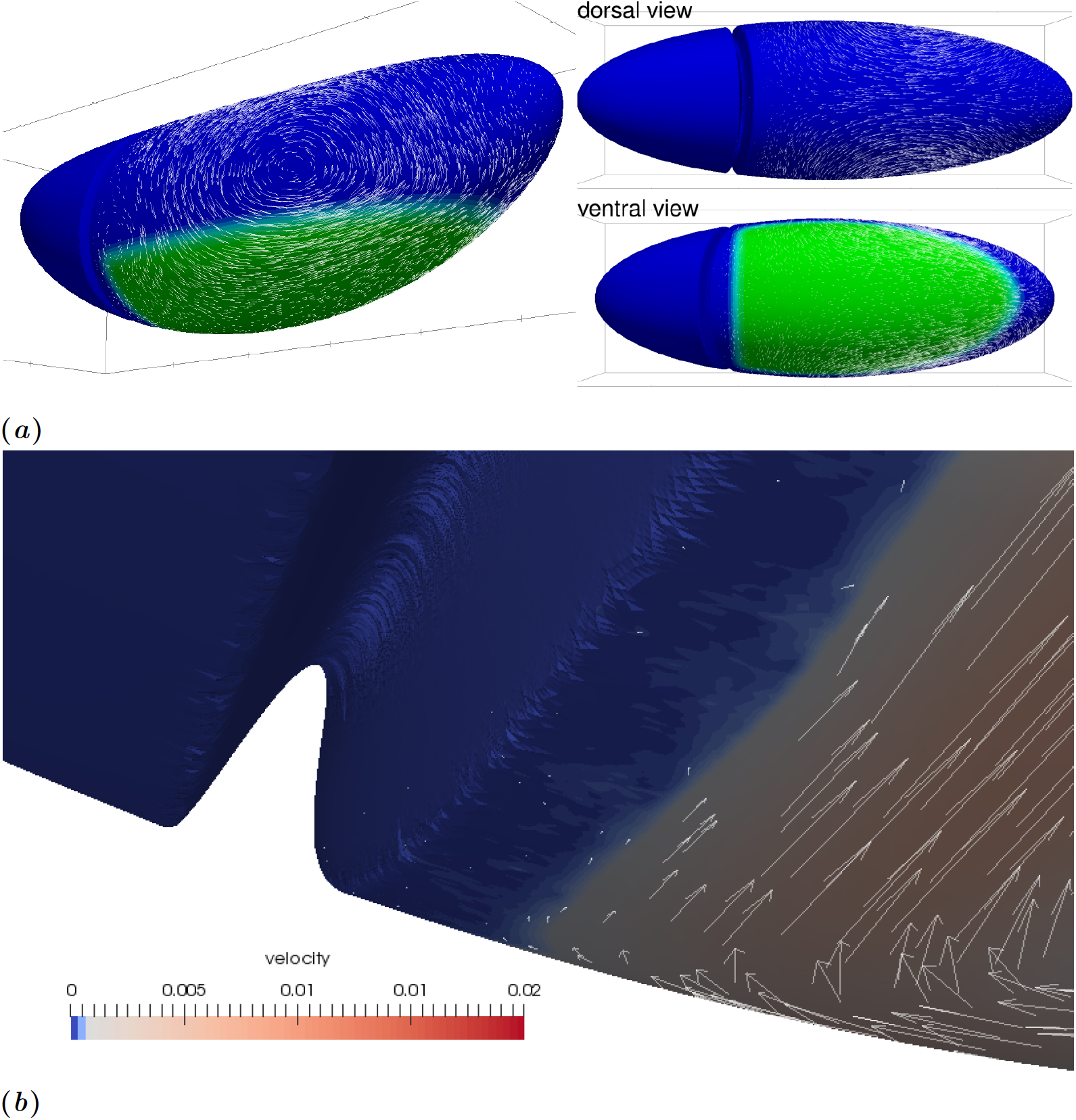
Flow field generated by planar-polarised myosin contractility in the GB. Parameters are *η*_b_*/η* = 10^3^, *c*_f_*/η* = 1*/R*, where *R* is the radius of a transverse cut of Γ, see Fig 1*d*. The GB region is defined as the ventral region posterior to cephalic furrow, and more ventral than a coronal plane *z*_max_ = −0.2*R*, see Fig 1. (*a*) Global view. The cephalic furrow is represented by a trough, the ventral furrow is not represented as we assume it to have sealed completely at the time corresponding to the simulations. Green, ventral region of the GB where we assume myosin planar-polarised contractility. White arrows, velocity vectors (arbitrary units, not all vectors calculated are represented). (*b*) Close-up of the region close to cephalic furrow and ventral midline. Every velocity vector calculated is represented, in arbitrary units 10 times larger than in panel (a).

In order to compare this flow qualitatively with the actual movements of epithelia during *Drosophila* germband extension, we have calculated the displacement of the centroid of cells *in vivo*. We have imaged a wildtype *Drosophila* embryo expressing plasma membrane associated GFP markers using light sheet microscopy (specifically mSPIM, multidirectional selective plane illumination microscopy) to observe the whole embryo volume throughout germband extension in four perpendicular views then reconstructed to four-dimensional movies [6]. Using custom built software (see S1 Text) we mapped apical cell trajectories on one lateral surface of the embryo. In Fig 1*a*, the displacement between two consecutive frames during the fast phase of GB extension is shown. Such *in toto* tracking permits an account of the tissue movement across the whole of the embryo simultaneously, compared to previous microscopy techniques which were restricted to the observation of a limited view field [9]. The global features of the flow predicted by our myosin-activity based simulations are also apparent in this tracking. There is an embryo-scale vortex on each lateral side of the trunk part of the embryo, with the same direction of rotation between simulations and observations. The morphogenetic flow is much reduced in the head region, in simulations this reduction is even stronger, and velocities in the head region are less than 5% of the maximum velocity. Discrepancies between the model prediction and the actual flow can be reduced when biophysically relevant mechanical parameters are adjusted, and when the PMG invagination effect is included: this is the objective of the next sections.

### Complementary role of planar-polarised myosin and posterior midgut invagination

It has been shown [6,7,14] that GB extension is not solely due to the action of planar-polarised myosin within the GB, but also to the pulling force that another morphogenetic movement causes, namely the invagination of endoderm in the posterior region, also called post-midgut (PMG) invagination.

Our present numerical approach does not allow us to simulate deformations of the surface in the normal direction, which would be necessary to simulate the PMG invagination process. However, it is possible to mimick the effect of PMG invagination on neighbouring tissues by simulating an in-plane isotropic contraction of a posterior patch of tissue, Fig 2*b*. Fig 4*e* shows that this does generate an extension of the GB area, although the deformation on the dorsal side is greater in this case. Combined with planar-polarised myosin action in the GB, Fig 4*b–d* shows that posterior contraction does modify the flow pattern substantially and provides a complementary cause of GB extension, consistently with the experimental studies cited above. In fact, because of the linearity of Eq (1)–(2), the superposition principle applies and the flows shown in Fig 4*b–d* can be written as the weighted sum of the flows with planar-polarised myosin only, Fig 4*a*, and posterior contraction only, Fig 4*e*. The location chosen for posterior contraction corresponds to the location of PMG in early GB extension, Fig 1*b*, rather than the stage for which we have tracked cell flow, Fig 1*a*. In spite of this, the agreement with the experimental observations, Fig 1*a*, is improved by the addition of posterior contraction. The flow within the posterior region itself is not relevant to compare, since the numerical method does not allow for the invagination in itself. In neighbouring tissues, posterior contractility modifies the predicted flow on the dorsal side, creating a posterior-ward flow in a small region close to the contractile patch, and reducing the flow rate in the rest of the dorsal side. In particular, the posterior contractility in Fig 4*b* leads to flow rates of similar relative magnitude to the ones observed experimentally in Fig 1*a*.

**Fig 4:**
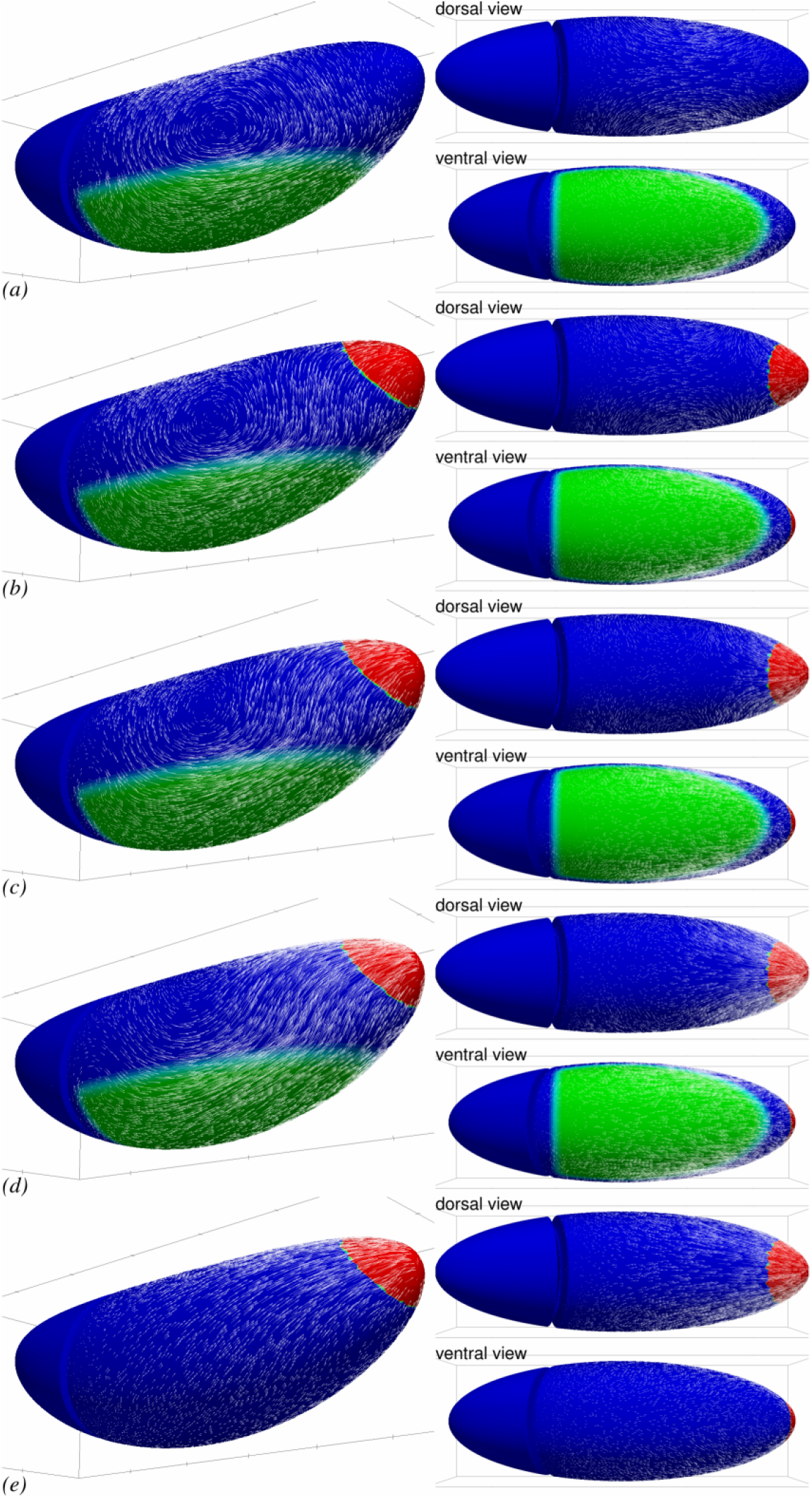
PMG invagination can contribute to GB extension. Parameters are the same as in Fig 3, reprinted in (*a*), but (*b*–*e*) another region (red) is actively contracting in an isotropic way, mimicking the effect of PMG invagination on neighbouring tissues. From (*b*) to (*d*), the isotropic PMG contraction intensity is doubled each time. (*e*), effect of isotropic PMG contraction in the absence of any myosin activity in the GB itself.

### Influence of the ventro-lateral patterning of myosin activation

Although *in toto* imaging of *Drosophila* embryos is now possible using SPIM to track the morphogenetic movements which our model attemps to predict [6,9,26], see also Fig 1*a*, a global cartography at the embryo scale of myosin localisation and activation during GB extension is is not available yet. The patterning of gene expression upstream of Myosin II planar-polarised recruitment, on the other hand, is accurately described [13]. We tested several configurations of the extent of the region of GB where myosin is anisotropically localised, Fig 5. We find that the global flows present a common aspect and GB extension is consistently obtained. However, some features vary, and in particular an anterior-wards backflow develops along the ventral midline when the myosin activation zone extends more laterally, Fig 5*b*. These backflows are attenuated or disappear when the PMG contraction is large enough, by superposition with flows in Fig 4*e*.

**Fig 5:**
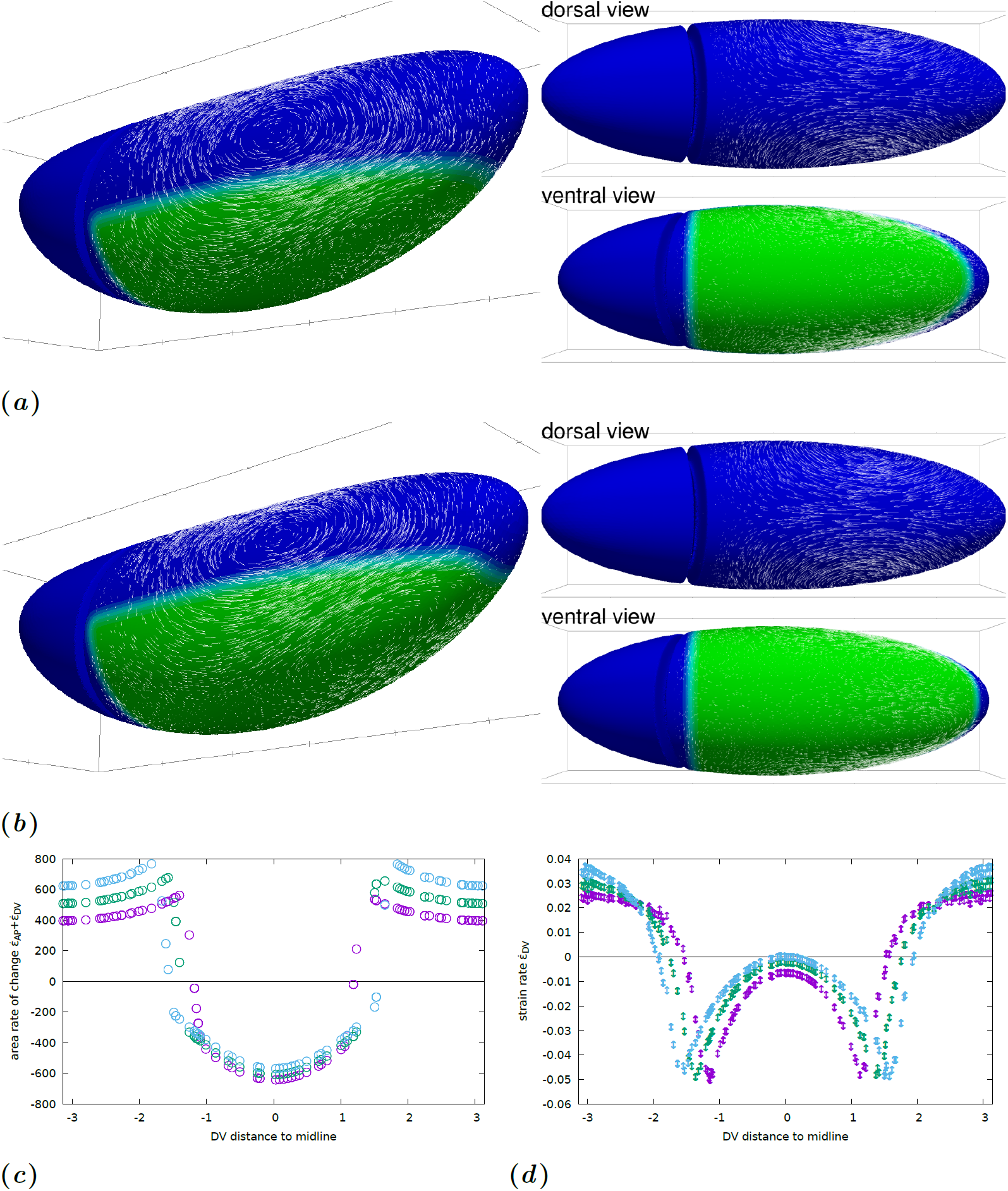
Influence of the patterning of myosin activation. (*a*–*b*) Flow field generated by planar-polarised myosin contractility in the GB. Parameters are the same as in Fig 3, but the region of planar-polarised myosin recruitment is larger: (*a*) *z*_max_ = 0, (*b*) *z*_max_ = 0.2*R* (compare to Fig 3 where *z*_max_ = *−*0.2*R*). The same phenomenology is observed, with a strong posterior-oriented GB extension. Close to the midline, the tissue is not extended and goes towards anterior (slightly for *z*_max_ = 0, significantly for *z*_max_ = 0.2*R*). (*c*) Rate of area change as a function of the DV distance to ventral midline (in units of *R*), along a transverse cut midway along AP (*x* = 0), for *z*_max_ = *−*0.2*R* (*purple* symbols), *z*_max_ = 0 (*green* symbols), and *z*_max_ = 0.2*R* (*cyan* symbols). In the region where myosin prestress is nonzero, area decreases, while it increases outside this region. (*d*) Rate of DV strain as a function of the DV distance to ventral midline (in units of *R*), along a transverse cut midway along AP (*x* = 0), for *z*_max_ = *−*0.2*R* (*purple* symbols), *z*_max_ = 0 (*green* symbols), and *z*_max_ = 0.2*R* (*cyan* symbols). Dorsally, there is a DV extension and ventrally a DV contraction. A peak of DV contraction is located at the boundary of the myosin activated region in each case.

In terms of the surface deformation of the epithelium, it is seen in Fig 5*c, d* that the flow is qualitatively the same although the location of the boundary of the myosin activated region directly changes the location of the strong gradients of strain rate. Within each region, the simulation reveals nontrivial variations of the strain rate. We note in particular strong peaks of deformation rate close to the boundaries of the myosin activated region (Fig 5*d*), which are not reported in *Drosophila*. These peaks are strongly reduced if the myosin prestress is assumed to change smoothly rather than abruptly at the boundary of the region, as will be seen below. Overall, the similarity of the flows observed from *z*_max_ = *−*0.2*R* (Fig 3) and *z*_max_ = 0 (Fig 5*a*) and their qualitative agreement with experimental observations (Fig 1*a*) is indicative that the system is robust with respect to the exact pattern of myosin activation.

### Influence of the choice of the mechanical parameters

Four parameters appear in Eq (3): the magnitude of myosin prestress *σ*_a_, the friction coefficient *c*_f_, and the two viscosities *η*_b_ and *η*. Two of these parameters, *σ*_a_ and *η*, set respectively the magnitude of stresses and velocities, leaving two free parameters: *η*_b_*/η*, which is nondimensional; and *c*_f_*/η*, which is the inverse of the hydrodynamic length Λ.

The ratio *η*_b_*/η* compares the bulk to the shear viscosity: if it is very large, then the flow will be nearly incompressible in surface, meaning that any surface element (and in particular, any cell) will conserve the same area through flow, and hence all deformations will be locally pure shear deformations. If the ratio is small, then the viscous cost of locally changing the apical area will be similar to that of pure shear, and, depending on the global force balance, area changes may dominate. Indeed, if *η*_b_*/η* is ^2^*/*3 (such that the Poisson ratio is zero), then a unixial load (such as a perfectly planar-polarised myosin action could be supposed to produce) will result in an area change only and no pure shear at all, or, in developmental biology terms, in convergence only and no extension. This effect is illustrated in Fig 6, where *η*_b_*/η* covers the range 10 to 10^3^. In the latter case, the negative DV strain rate (convergence) exactly balances the positive AP strain rate (extension), whereas in the former case, convergence strongly dominates. The strain rates in all cases are not uniform across the ventral side depending on the AP position, with a marked decrease of the pure shear in the central part of the GB, however other factors are seen to affect this spatial distribution in the rest of the paper. Between *η*_b_*/η* = 10 and *η*_b_*/η* = 100, there is a switch from area-reduction dominated flow (area reduction rate 4 times AP strain rate) to a shear-dominated flow (area reduction rate of the same order as AP strain rate). Experimentally in *Drosophila* embryos, the deformations are not limited to pure shear but involve some area change [14]. By comparison of actual tissue flows in Fig 1*a* with our numerical result, the order of magnitude of *η*_b_*/η* can be expected to be of the order of 10 to 100.

**Fig 6:**
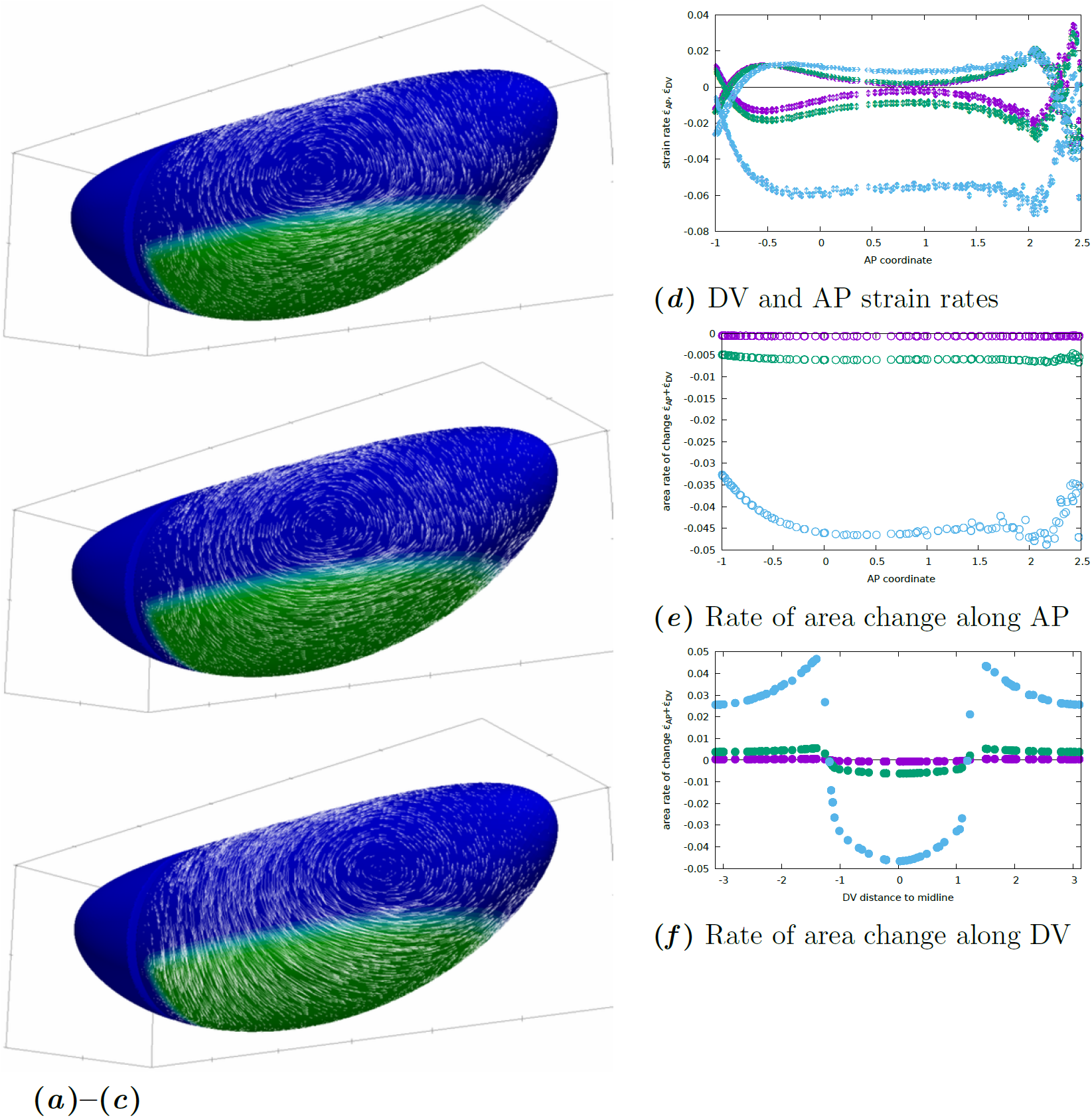
Influence of the bulk viscosity on the convergence and extension of GB. (*a*–*c*) Flows calculated for (*a*) *η*_b_*/η* = 10^3^, (*b*) *η*_b_*/η* = 10^2^, (*c*) *η*_b_*/η* = 10. While the posterior-wards flow at the GB posterior end is similar, the lateral flow is strongly affected with a much larger ventral-wards convergent flow along DV when the bulk viscosity is reduced to 10 (whereas no qualitative change is seen between 10^3^ and 10^2^). The position of the vortex centre is also much modified for low bulk viscosity. (*d*) Rates of strain in the DV (↕ symbols) and AP (*↔* symbols) directions for the three choices of *η*_b_*/η* (*purple*, 10^3^, *green*, 10^2^, *cyan*, 10), as a function of AP coordinate *x* (in units of *R*) close to the midline (*y* = 0.2*R*). In the GB, DV rate is negative (convergence) and AP rate positive (extension). The DV rate of strain is increasingly negative for low bulk viscosity, indicating a stronger convergent flow, while the AP rate increases much less, indicating little change in the rate of GB extension. Posterior to the GB (AP coordinate *x*;: 2*R*), the DV and AP strain rate values ramp and invert their sign, indicating that the direction of elongation swaps from AP to DV, which corresponds to the splayed velocity vectors seen at the posterior limit of the GB e.g. in panel *a*. Anterior to the GB (AP coordinate *x ≃ −R*), the same effect is observed due to the obstacle of the cephalic furrow. (*e*) Rate of area change for the same choices of *η*_b_*/η*, confirming that area decreases much more for lower bulk viscosity in the GB region. (*f*) Rate of area change for the same choices of *η*_b_*/η* as a function of the DV distance (in units of *R*) to ventral midline, along a transverse cut midway along AP (*x* = 0). When *η*_b_*/η* is sufficiently small to allow area variations, the GB region exhibits area reduction and dorsal region area increase.

The hydrodynamic length Λ = *η/c*_f_ is the characteristic length within which shear stress will be transmitted within the actomyosin. Beyond this length, friction with the exterior will balance the internal stress. It thus indicates the length over which the effects of a local force are felt. We tested the effect of a hydrodynamic length either much larger, much smaller or comparable to the size of the embryo (*R*, the radius of a transverse section, Fig 1). The results shown in Fig 7 show how the action of myosin gives rise to a more local flow pattern when the hydrodynamic length is small. This localisation is around the areas in which there is a *gradient* of myosin activity. In areas of uniform myosin activity, the resulting effect is a uniform tension (see a similar effect in models of cells plated on a substrate, e.g. [40, Fig. 3*b*]). It can be seen on Fig 7*e,f* that indeed the strain variations are more abrupt when hydrodynamic length is small, whereas the long range interactions allowed by a very large hydrodynamic length give rise to an embryo-scale flow. On the whole, flows which reproduce experimental observations better are obtained when Λ is of order 1 or more, that is, the hydrodynamic length is comparable to or larger than the GB width in DV. Indeed, there is no strong localisation of the flow features in our exeperimental results, Fig 1*a*. This is consistent with the order of magnitude of 10 to 100 *µ*m found by laser ablation in other systems [43].

**Fig 7:**
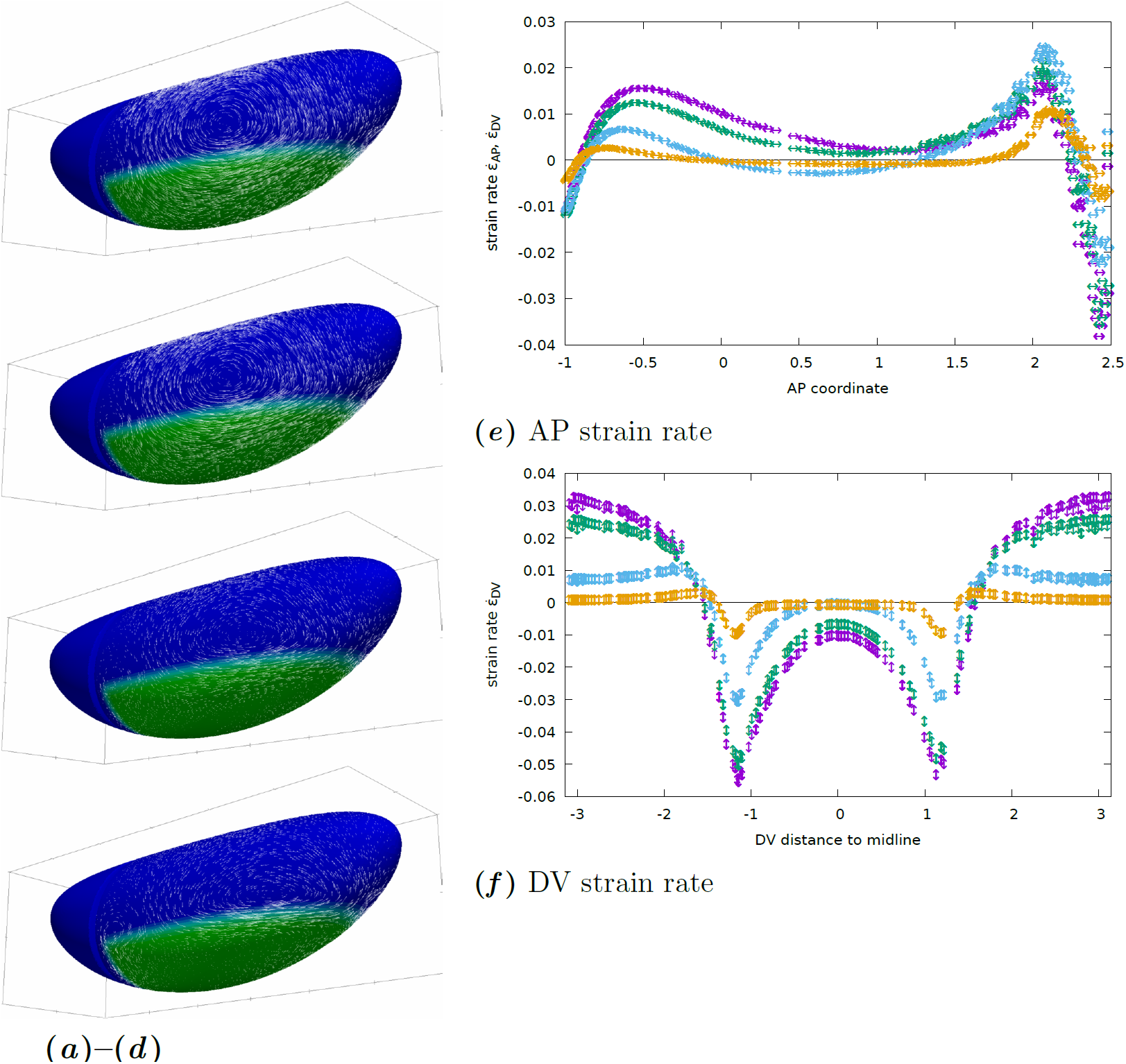
Friction with vitelline membrane and/or cytosol and yolk modifies the flow pattern. (*a*–*d*) Global flow pattern. Parameters are the same as in Fig 3, but the hydrodynamic length varies between Λ = 10*R* (*a*), Λ = *R* (*b*, reprinted from Fig 3), Λ = *R/*10 (*c*), and Λ = *R/*100 (*d*). In all cases the GB extends posteriorly. In the cases of small hydrodynamic length, convergence and extension flow occur mostly in regions where there is a gradient of contractility. Overall, friction renders the effect of actomyosin activity more local to regions where they exhibit a variation, hence vortex structures are more localised next to these regions with higher friction and have less influence in regions of uniform actomyosin activity (dorsally or close to ventral midline e.g.) (*e*) Rates of strain in the AP direction for the four choices of Λ*/R* (*purple*, 10*R*, *green*, 1, *cyan*, 1*/*10 and *orange*, 1*/*100), as a function of AP coordinate *x* (in units of *R*) close to the midline (*y* = 0.2*R*). The overall magnitude of strain decreases with increased friction, as an increasing part of the energy provided by myosin activity needs to overcome friction in addition to deforming the cell apices. The rate of strain is uniformly positive (elongation) only when Λ is close to unity or smaller, else a region of shortening appears in the central part of GB. (Note that since area change is close to zero, the DV strain value is very close and opposite to the value of AP strain everywhere.) (*f*) Rate of strain in the DV direction for the four choices of Λ, as a function of AP coordinate *x* (in units of *R*) close to the midline (*y* = 0.2*R*). DV rate of strain always peaks close to the boundary of the GB area where myosin is active, in relative terms the peak is more pronounced for large frictions. Dorsally, there is always a positive DV rate of strain, indicating a DV elongation due to the pull of the neighbouring converging GB. This is matched with an AP shortening of a similar magnitude. Ventrally, the negative rate of strain (indicating convergence) is observed to decay when the hydrodynamic length becomes small, in that case the DV narrowing is limited to a narrow band at the DV edge of the GB. This localisation effect of small hydrodynamic length is also seen for the dorsal DV elongation, but to lesser extent.

### The cephalic furrow acts as a guide for morphogenetic movements

In simulations, Fig 3, it is seen that the flow follows the cephalic furrow in a parallel way. Thus we wondered whether the presence of the CF could be important for the flow pattern observed. To test this, we performed the same simulation on two different meshes, one featuring the CF and the other without it. In the absence of a CF, the flow at the anterior boundary of the GB does not deviate laterally but continues towards the anterior, Fig 8*e*. The flow field is thus much more symmetrical than in when the CF is included in the model, Fig 8*b*. From the mechanics, we expect that if both the geometry of the embryo and the myosin localisation patterns are symmetric, then the flow will be symmetric too (see Fig 8*d* for a verification of this). In real embryos, two sources of asymmetry arise: the localisation of the posterior boundary of the region of planar-polarised myosin recruitment pattern, and the invagination in the posterior midgut. Blocking it however does not completely suppress the posterior-ward extension of GB [7]. Our results suggest that another asymmetry could originate from the geometry of the embryo, with the CF acting as a barrier resisting flow towards the anterior. Indeed, if one introduces this geometric feature at one end of the otherwise perfectly symmetrical embryo, the flow is strongly asymmetric towards the posterior end, Fig 8*a*. In experimental embryos, the local influence of the CF on the flow is visible in tracked *in toto* images, Fig 1*a*, where a reduction of the magnitude of the velocity is seen in the row of cells posterior to the CF. The numerical simulations bring the additional understanding that, in addition to this local effect, this mechanical feature has a global influence on the morphogenetic flow and orients GB extension towards the posterior.

**Fig 8:**
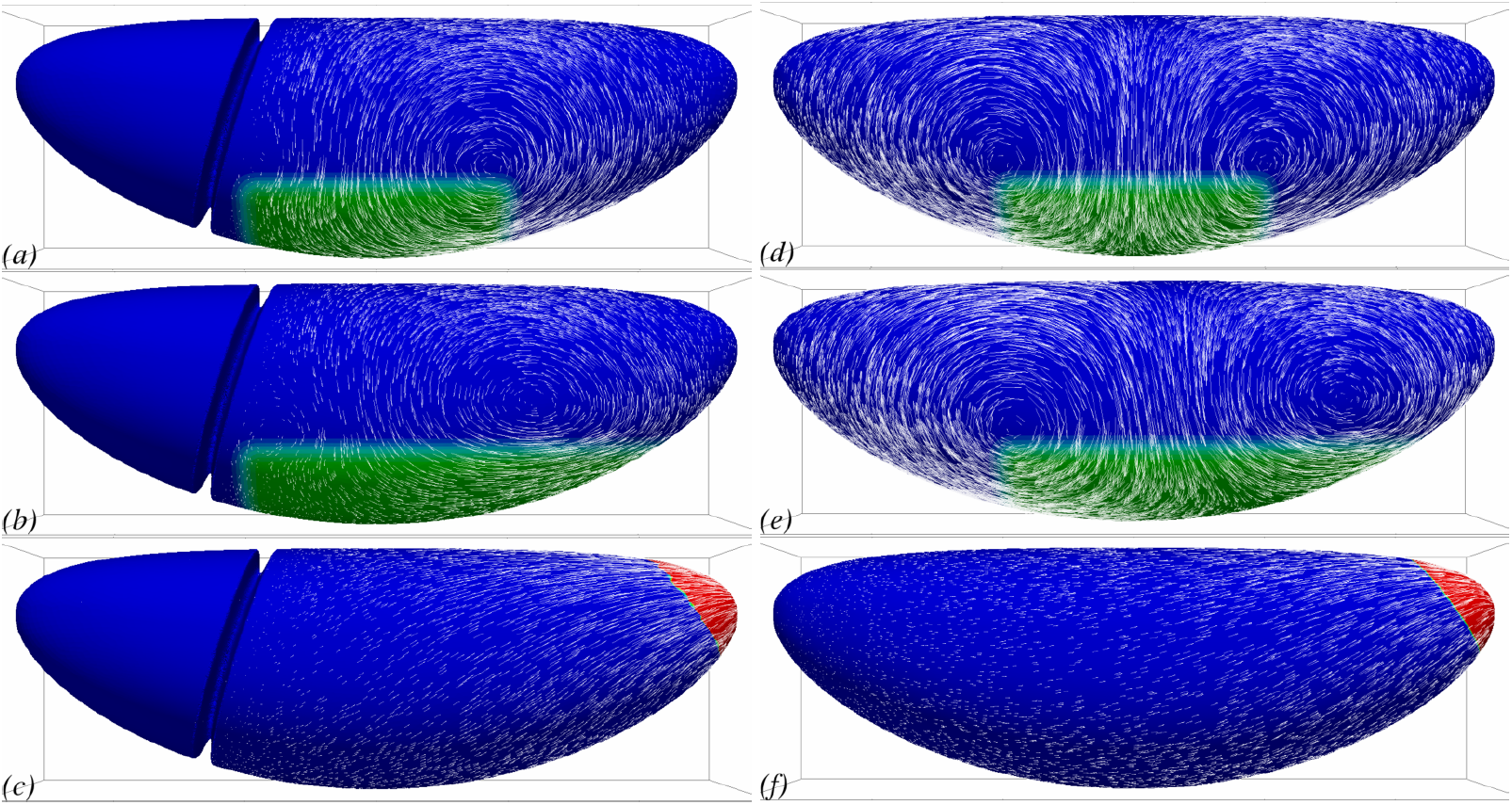
Cephalic furrow (CF) can guide GB extension to be mostly posterior-wards. Lateral view of flow fields generated by myosin contractility in the presence (*a*–*c*) or absence (*d* –*f*) of a cephalic furrow. (*a*,*d*) With a hypothetical symmetric planar-polarised myosin activity, the presence of CF orients the flow towards the posterior whereas it is perfectly symmetric in its absence. (*b*,*e*) With a realistic asymmetric planar-polarised myosin activity, the presence of CF still has a major role in orienting the flow the towards posterior. Although the asymmetric myosin patterning induces a asymmetric flow in the absence of the CF, the flow is not biased towards the posterior. (*c*,*f*) The flow created by PMG invagination is much less sensitive to the presence of CF.

### Simulation of wild-type GB extension

Using the qualitative study of the influence of each parameter (*η*_b_*/η*, *c*_f_*/η*) and of the patterning of myosin activation, we propose a choice of simulation settings which can be expected to reproduce the features of GB extension flows. Indeed, the equations being linear, the superposition principle guarantees that the effect of each of these causes simply add up. In Fig 9*a*, the magnitude of PMG strain rate is as in Fig 4*b*, *η*_b_*/η* is chosen as in Fig 6*b*, the hydrodynamic friction as in Fig 7*b*. The myosin prestress pattern is close to the one in Fig 5*a*, but with a graded decrease of prestress when reaching the boundary of myosin activated region. Fig 9*b–e* show the rate of strain along lines of interest on the surface. Compared to similar plots where myosin prestress is not graded close to the boundary, they have the same global features but no peaks close to the myosin activation boundary: compare e.g. Fig 5*d* and Fig 9*e*. This is indeed due to the gradation of myosin prestress, since a simulation where only this parameter is altered restores the peaks (Supp. Fig. S1).

**Fig 9:**
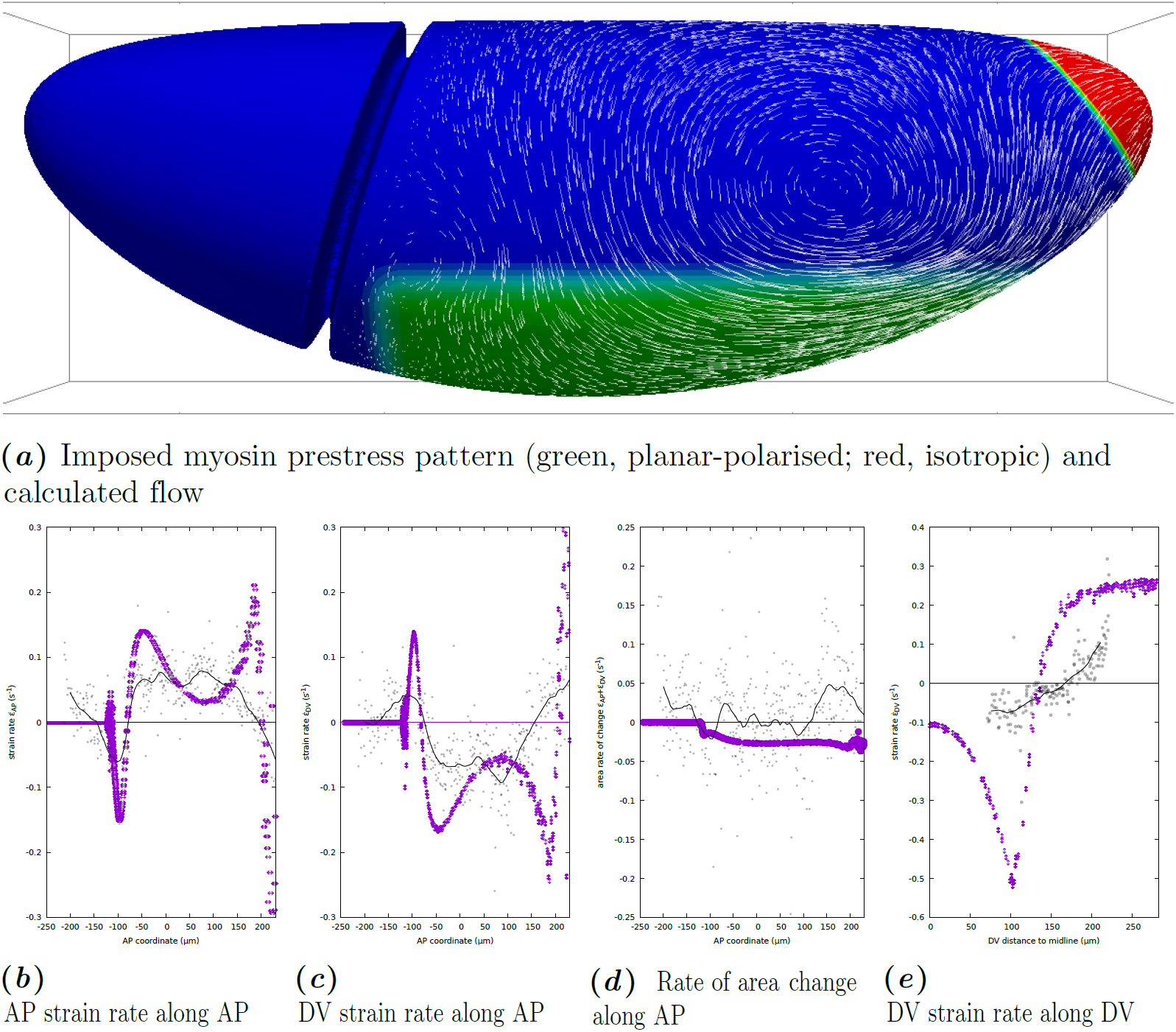
Simulation of GB extension with proposed parameters for the WT *Drosophila* and comparison with an example of real data. (*a*) Flow calculated for a proposed choice of parameters *η*_b_ = 10^2^*η*, *c*_f_*/η* = 1*/R*, and a proposed choice of myosin prestress distribution and polarisation, as shown in colour code. The intensity of the prestress is graded at the boundary of the myosin-activated region. (*b*–*e*) Rates of strain predicted by the simulation (*purple symbols*) along specific AP or DV-oriented lines, and measured from the example of tracked data presented in Fig 1*a* (tissue strain rate calculated at cell centroids, gray dots; LOESS regression with 25-micron window, black curve). (*b*) Rate of strain in the AP direction as a function of AP coordinate *x* close to the midline (*y* = 0.2*R*). Rate of strain in the DV direction as a function of AP coordinate *x* close to the midline (*y* = 0.2*R*). (*d*) Rate of area change as a function of AP coordinate *x* close to the midline (*y* = 0.2*R*). (*e*) Rate of strain in the DV direction as a function of the DV distance to ventral midline, along a transverse cut midway along AP (*x* = 0).

In Fig 9*b–e*, we also plot real data of tissue strain based on the velocity of cell centroids on the surface of an embryo at one time point of GB extension, shown in Fig 1*a* [15] (see also Methods). In order to do this comparison, we use the known value *R* = 90 *µ*m, and adjust manually the time unit of the simulations (one scaling parameter). It is seen that the spatial dependence of each quantity of interest follow similar patterns of variations in the simulation and the real data.

Along the AP direction, Fig 9*b–c*, there is a systematic shift of the position of the first peak value, which is located immediately posterior of the CF (*x* = *−*100 *µ*m) in simulations, and straddles it in real data. This may be due to the incompleteness of the mechanical model of the CF, which in simulations appears as an immoveable fold, while in reality this fold has some limited movement, and deepens over time during GB extension as cell apices enter it [44]. In spite of these differences, the magnitude and shape of the peaks of strain associated with the CF are accurately reproduced in the simulations.

Further into the GB, *−*25 *µ*m≲ *x* ≳ 175 *µ*m, simulations and real data exhibit a plateau of negative DV strain (convergence), Fig 9*c*, and positive AP strain (extension), Fig 9*b*, with little area variations, Fig 9*d*. as already discussed in Fig 6*d*. At both the posterior and anterior (CF) end of the GB, there are regions in which the sign of the rate of strain invert. This is consistently observed in experiments and simulations. In these regions, there is thus a local convergence-extension process in the direction orthogonal to the one in the GB. In real data, part of the convergence in AP is due to the loss of tissue as it goes into the CF and deepens it [44], and there is a corresponding peak of area decrease. In simulations, the AP convergence at the two ends of the GB is driven by the push from the AP extending GB, and the DV extension to the resistance to area changes, see also Fig 6*d*.

Along DV, Fig 9*e*, the current tracking technique of 3D SPIM data offers a range limited to ca. 90 degrees in DV. Over this range, the agreement in terms of the magnitude of the DV strain is rather good, but its rate of variation along the DV direction is not accurately predicted. This may be linked with an inacurate hypothesis on the gradation of myosin prestress close to the boundary of the GB, since Supp. Fig. S1*e* exhibits a very different slope from the simulation in Fig 9. Dorsally, simulations and experiments agree on the existence of an AP convergence and a DV extension. It is consistent with observations in [9] that cells in this dorsal region extend in the DV direction while not rearranging and incurring little area change.

Overall, there is a good agreement in qualitative features in strain rates using the parameters that have been chosen based on flow features in the previous sections. That the latter agreement would lead to the former is not trivial, and Fig 6*d-f*, 7*e*,*f* show that similar flows can exhibit different features in terms of strain rate.

## Discussion

### Predictability of GB extension flow from myosin distribution

Our simulations are based on the knowledge of the embryo geometry, which includes the global shape of the continuum formed by epithelial cell apices and also the fold formed by the cephalic furrow (CF), and an assumption of the pattern of myosin expression over this surface and its planar polarisation. A mechanical model (Eq (3)) then allows us to predict the stress and flow field over the surface of the whole embryo, using a novel numerical technique that can solve the equations on the embryo surface in three-dimensions [25]. The mechanical model is based on a liquid-like constitutive relation for the actomyosin of the apical continuum, following [21,39]. We only obtain a snapshot of the flow field that corresponds to a given myosin activation pattern. In the course of GB extension, this pattern is transported by the flow [13], and the flow field evolves in its details while preserving its global features [9]. Because there is no time evolution term in our model, Eq (3), there is a unique instantaneous flow field for a given pattern of myosin prestress ***σ*****_a_**. At a given time point, the results (Fig 9) compare well with corresponding observations of the morphogenetic movements of GB extension Fig 1*a*, which we track over the three-dimensional surface of the embryo using *in toto* imaging and tracking software [15]. This agreement suggests that the model captures the essential balance of stresses originating from the contractility of actomyosin. In order to go further into model validation, the next step will be to apply the model to measured embryo shape and myosin localisation patterns, which could in the future be obtained by *in toto* embryo imaging. This would allow the model to be validated by testing its ability to predict the flow that corresponds to each successive myosin localisation pattern and CF position in the course of GB extension.

We note that in order to obtain a good agreement of model predictions with observations in terms of strain, we have had to use a graded intensity of myosin prestress at the boundary of the planar-polarised actomyosin region, Fig 9, rather than use a sharp change, Supp. Fig. S1. It remains to be shown whether such a gradation of planar-polarised actomyosin can be observed experimentally. The DV patterning of myosin localisation, although important for the precise localisation of flow features, is not crucial to achieve a convergence-extension motion with the correct orientation, Fig 5. For a planar-polarised actomyosin region extending far laterally however, our simulations predict backflows close to the midline. Such backflows are not observed experimentally, although mutants such as *torsolike* that present planar-polarised myosin but not PMG invagination form ectopic folds in the germband [6], which could be due to a buckling phenomenon. The mechanical role of the invaginated mesoderm may have to be taken into account in order to improve our modelling of the midline region, Fig 1*d*.

In addition to the actomosin patterning, two mechanical parameters govern the flow we predict, the ratio of apical surface resistance to compression relative to its resistance to shear, and the hydrodynamic length. Both have a more general biophysical relevance (see e.g. [43]). The material parameters of the cortical actomyosin in *Drosophila* are not known, and quantifying them experimentally is challenging [39], since this early step of development requires the presence of the rigid vitelline membrane that surrounds the embryo and prevents direct mechanical measurements from the exterior. Magnetic tweezers have been used [45], but the magnetic particles were not directly associated with the actomyosin cortex and thus measured other mechanical properties than those required by our model. Laser cuts can provide valuable information on the evolution of tension [46] or its anisotropy [6]. By comparison with a model of subcellular actomyosin, its material parameters can be obtained (medial actomyosin relaxation time, shear viscosity and friction coefficient) [43]. These values correspond to a subcellular system while we focus on the tissue scale, which may lead to quantitative differences with our case. Indeed, although the model in [43] is formally identical to Eq (1)–(2), their friction coefficient includes the friction cost of movements of the actomyosin cortex relative to the cell membrane, whereas both apical membranes and actomyosin cortices flow in GB extension. In spite of this, the hydrodynamic lengths of 14 to 80 *µ*m found for actomyosin cortex of single-cell *C. elegans* embryos and actomyosin ring of gastrulating zebrafish respectively in [43]. The ratio of bulk to shear viscosity *η*_b_*/η* that we retain is much larger than the ratio 3 that could be expected from a simple 3D-isotropic modelling of actomyosin [43]. This is important in the model in order for the DV stresses to lead to AP extension as well as DV convergence, since it is the bulk viscosity that couples these effects, Fig 2*d*. The origin of this large resistance to area variations could be due to the combination of the volume constraint of each cell and of a basolateral regulation of its height, as has been hypothesised in cell-based models [30]. Alternatively, active processes may regulate the apical actomyosin cortex thickness and density.

The method can account for the fine geometrical detail of the embryo apical surface, such as the CF. It is observed experimentally that during GB extension, the cell displacements are much less in the head region than in the trunk, Fig 1*a*. However, there is no report of a specificity of the cytoskeleton of cells in the head at this stage that could account for a locally enhanced stiffness. Our simulations lead us to propose that the presence of the CF may be the cause for these smaller displacements. Geometric features such as the CF are important for the mechanical equilibrium, since forces are transmitted directionally and any curvature will modify the equilibrium. We show that the CF modifies strongly the flow we predict and acts as a barrier for deformations, Fig 8. This geometrical feature thus guides the convergence–extension flow towards the posterior end of the embryo.

### A mechanical scenario for GB extension

Our results confirm that either of the two mechanisms whose elimination was seen to correlate with a reduction of GB extension [6,7,11,14,42] can be the direct mechanical cause of a flow towards the posterior in the GB. The results in Fig 4 indicate that the GB extends under the effect of the anisotropic prestress of GB planar-polarised actomyosin, but also under the effect of the pull from the invaginating PMG. However, the precise flow patterns differ in these different cases. An important and obvious next step is therefore to obtain experimentally the distribution and polarisation of myosin on the entire embryo surface across time, for example using SPIM, for wildtype and different mutants, which should allow us to quantify the parameters and test the predictive power of the model. In the interval, our theoretical work already sheds light on the fundamental mechanisms at play and how they integrate in the complex 3D geometry of the embryo to yield the morphogenetic events that are observed.

Based on our simulations and the timings reported in the literature, we can indeed articulate a mechanical scenario for GB extension. The endoderm contraction that leads to PMG invagination, the first event correlated with GB extension [6, Fig. 4*B*], starts several minutes before the onset of GB extension can be detected. In *Kruppel* mutants, for which the planar polarisation of myosin in GB is deficient, GB still extends [14], this is mainly due to PMG invagination [6]. The corresponding simulation is shown in Fig 4*e*, PMG contraction generates a flow towards the posterior that does extend the posterior half of the germband but decays rapidly in space. The presence of the cephalic furrow for this extension does not have a strong influence there, see Fig 8*f*. Thereafter, from shortly before the onset and in the course of GB extension, myosin becomes increasingly planar-polarised [19]. The direct consequence of this is a lateral flow from dorsal to ventral, causing *convergence*, that is, a negative rate of strain of GB along the DV direction. Due to a rather large value of *η*_b_, which corresponds to the in-plane compressibility viscosity of actomyosin, this causes GB extension, see Fig 6. Experimental evidence of such an in-plane low compressibility exists [7,14], although it is not clear whether this is a passive mechanical property of the actomyosin cortex or an active one [7,41]. In the absence of the cephalic furrow, Fig 8*e*, this extension occurs evenly in the anterior and posterior directions, in presence of the cephalic furrow, the viscous cost of flowing around the posterior end is much less than the cost of flowing into the furrow, and planar polarisation driven GB extension is biased towards posterior, even if the contribution of PMG invagination is not accounted for, Fig 3.

### Cell intercalation may not be necessary for convergence and extension of planar-polarised tissue

The two prominent features in which the existence of cell-cell junctions are important in GB extension in wildtype (WT) *Drosophila* are the planar-polarised recruitment of myosin, which preferentially enriches DV-oriented junctions, and the medio-lateral cell intercalations [11,13,42]. Because the DV-oriented junctions present both the characteristics of being enriched in myosin and of undergoing shrinkage to lead to intercalation, these two effects have so far been studied in association. However, some mutants such as *eve* that lack planar polarisation of myosin can still exhibit some cell intercalation, although to a much lesser proportion than cell shape changes [14]. Here, using our modelling approach we can envision the reverse case of studying convergence-extension due to planar-polarised myosin activity but without explicit cell intercalation. We show that the anisotropy of planar-polarised myosin activity is sufficient to explain convergence-extension, without the need for an intercalation mechanism.

We conclude that a mechanical model that does not involve individual cells but only a continuum standing for the apical actomyosin connected from cell to cell by apical junctions can produce a flow with strong similarities to GB extension. In vivo, cellularisation is of course important for the planar polarisation of myosin in the GB, since the DV-oriented cell-cell junctions are the location at which oriented actomyosin structures assemble. It is observed that acellular embryos do not exhibit myosin polarisation [6]. In the model however, it is possible to introduce planar polarisation independently of cellularisation. This is what we do when defining an anisotropic myosin prestress ***σ*****_a_**. The fact that no further account of the cellularisation of the embryo is necessary in the model suggests that at the tissue scale, one can address morphogenetic questions by considering ensemble displacements. In this approach, the effect of cell intercalation, which is governed by planar-polarised junctional myosin, is thus not directly taken into account, but rather encapsulated in a global tissue strain rate and its associate viscosities *η* and *η*_b_, which also include the cell deformation [15], Fig 2*f*.

This tissue strain rate 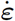 and the corresponding tissue-scale tension ***σ*** are related by the constitutive relation, Eq (2), which includes the contractility term ***σ*****_a_** resulting from planar-polarised myosin activity, and is thus the only term bearing a trace of the embryo’s cellularised organisation. The respective values taken by 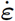 and ***σ*** locally depend on the mechanical balance, i.e. both the local myosin activity and the tension transmitted by neighbouring tissue, see Fig 2*d*. In the context of convergence-extension caused partly by invagination of the PMG, [14], it has been proposed that cell intercalation could relax the stress by allowing cell shape changes in the GB as it is extended by an extrinsic force. Here we propose that the crucial role of cellularisation in planar-polarised tissue convergence-extension lies in the mechanism through which oriented contractile actomyosin junctions are created, and not in the subsequent tissue deformation. This suggests that if planar-polarisation of actomyosin could be achieved through some other mechanism (as is the case in the *Drosophila* tracheal tubule [23]), convergence and extension could still be obtained in an acellular tissue, thus in the absence of cell intercalation. In cellularised tissue, an important role of cell intercalation could thus be the relaxation of the cell strain generated by the convergence and extension process, as was already suggested [14].

### Geometry-governed mechanical balance as a messenger in early morphogenesis

Planar-polarised myosin in the GB is known to generate a global flow at the surface of the embryo. Our simulations show that the global flow which is generated by such mechanical activity is dependent on the pre-existing geometry of the embryo, such as the presence or absence of the cephalic furrow. Thus, a prior morphogenetic movement such as cephalic furrow formation can affect further movements via mechanical interactions only.

This “messaging” proceeds via the establishment of a different mechanical balance depending on the geometry of the embryo, rather than the diffusion of a morphogen, e.g. [47]. In the early embryo, the distance over which these forces are transmitted is likely to be much larger than in later organisms, as there is no extra-cellular matrix structure that will relieve actomyosin from part of the stress. Indeed, we find that the hydrodynamic length is likely to be at least the width of the GB, consistent with laser ablation results [43], which implies direct mechanical interaction at this scale.

This mechanical messaging behaves differently from biochemical messaging. Its speed of propagation is the speed of sound in the force-bearing structure, here, the actomyosin. It does not propagate in an isotropic way but in a more complex directional one, and contains directional information. Regions of interest within the embryo should thus not be treated as isolated systems, since a distant geometric property of the embryo can have a direct impact on the mechanical stress felt locally when intrinsic forces are being generated.

This prompts further development of computational tools such as the one we present. Tangential flows on curved surfaces are also observed in other epithelia (such as follicular epithelium of *Drosophila* ovaries), but is also relevant to cortical flows in single cells, prior to mitotic cleavage for example. Mechanical approaches of flat epithelia have shed light on many aspects of tissue growth and dynamics [19,48–50], in particular at the scale of a few cells, which is the relevant one for cell rearrangements. At the other end of the spectrum of tissue dynamics, 3D phenomenological models of shape changes during ventral furrow formation have been proposed [30]. Here we propose a first step in bridging the gap between these approaches, with the objective to be able to address complex morphogenetic events in their actual geometry, and thus to fully account for the influence of current morphology on the mechanical balance that leads to further morphogenetic movements.

## Acknowledgements

JE thanks Alexandre Kabla, Alexander Fletcher and Jonathan Fouchard for fruitful discussions. All the computations presented in this paper were performed using the Cactus platform of the CIMENT infrastructure (https://ciment.ujf-grenoble.fr), which is supported by Région Rhône-Alpes (GRANT CPER07-13 CIRA:http://www.ci-ra.org). Authors would like to thank Philippe Beys who manages the platform. Authors thank Région Rhône-Alpes (CIBLE and IXXI, all authors; CMIRA, JE), MD thanks Malian government and French embassy in Bamako”Bourse d’Excellences” programme, LIPHY and LJK (CNRS and Univ. Grenoble Alpes) for financial support. MD and JE thank ANR-12-BS09-0020-01 “Transmig” and ANR-11-LABX-0030”Tec21”, and are members of GDR 3570 *MécaBio* and GDR 3070 *CellTiss* of CNRS. JE thanks the Isaac Newton Institute for Mathematical Sciences for its hospitality during the programme *Coupling Geometric PDEs with Physics for Cell Morphology, Motility and Pattern Formation* supported by EPSRC Grant Number EP/K032208/1.

## S1 Text – Additional Materials & Methods

## Lagrange multiplier approach for tangential flows

We present a generic finite element approach allowing to address flows of finite surface-compressibility *η*_b_. We approximate the solution of vectorial equations set on a curved surface of 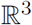. This also offers a framework for future extensions of the method to non-Newtonian fluids.

Eq (3) must be solved for velocities ***v*** tangential to the surface Γ, which corresponds to the continuum formed by the apical actomyosin cortices of cells and adherens junctions. In terms of solution spaces, this constraint can be written as 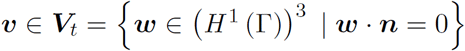 where (*H*^1^ (Γ))^3^ is the set of vector-valued functions defined on Γ whose differential is square-integrable, and ***n*** is the outer normal to Γ. Constraints such as the tangentiality of the solution can be implemented either by defining finite element spaces that can satisfy the constraint (see e.g. the work by [S1]) or by introducing a mixed finite element approximation following e.g. [S2]. Here we opt for the latter solution, which allows us to discretise vector fields on the surface in the Cartesian system of coordinates rather than a curvilinear one.

**Fig S1:**
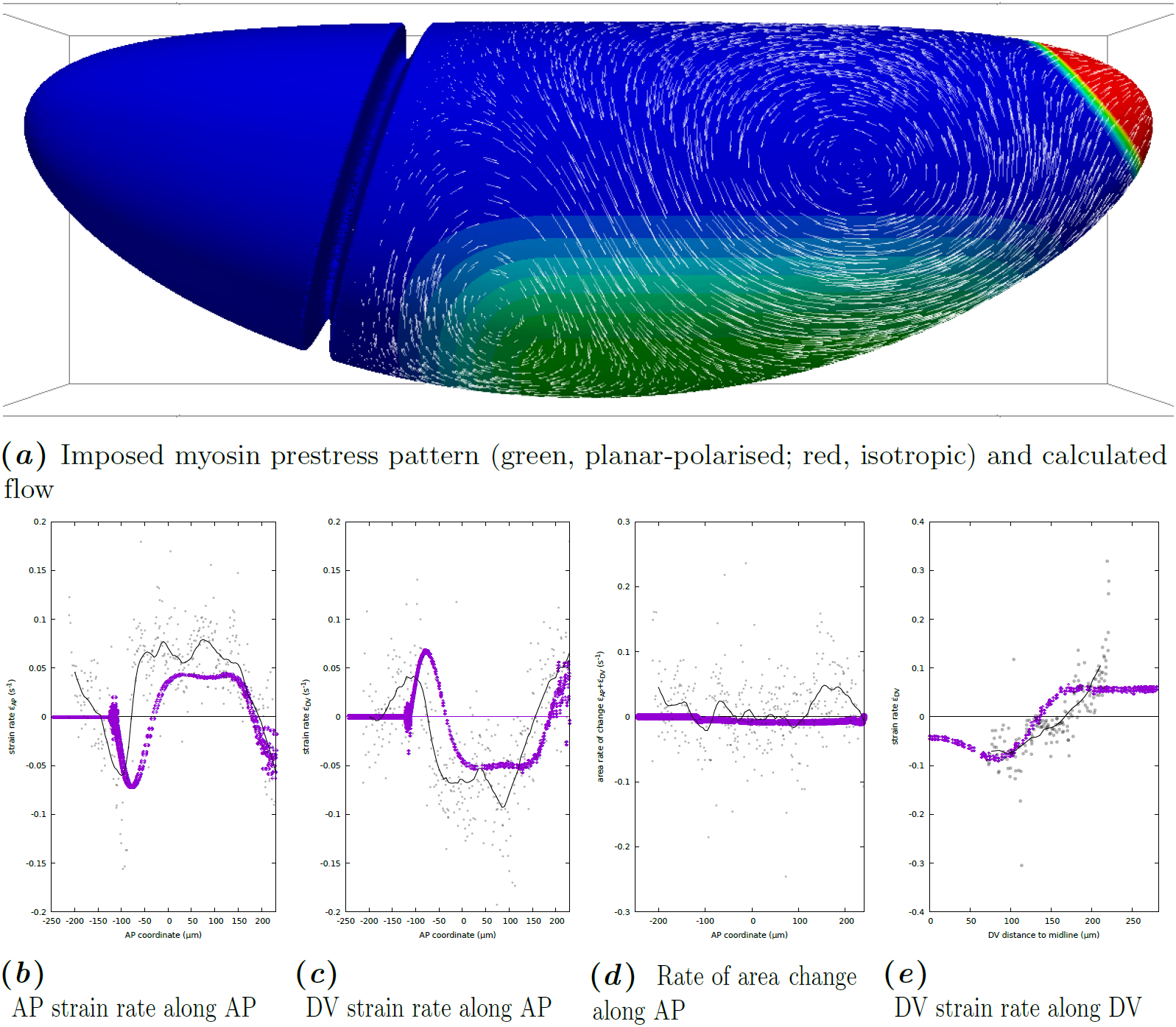
Simulation of GB extension with proposed parameters for the WT *Drosophila* but without graded myosin prestress (to the difference of Fig 9) and comparison with an example of real data. (*a*) Flow calculated for a proposed choice of parameters *η*_b_ = 10^2^*η*, *c*_f_*/η* = 1*/R* in colour code, and a proposed choice of myosin prestress distribution and polarisation, as shown. The intensity of the prestress is graded at the boundary of the myosin-activated region. (*b*–*e*) Rates of strain predicted by the simulation (blue symbols) along specific AP or DV-oriented lines, and measured from the example of tracked data presented in Fig 1*a* (individual cell rate of strain, gray dots; LOESS regression with 25-micron window, black curve). (*b*) Rate of strain in the AP direction as a function of AP coordinate *x* close to the midline (*y* = 0.2*R*). (*c*) Rate of strain in the DV direction as a function of AP coordinate *x* close to the midline (*y* = 0.2*R*). (*d*) Rate of area change as a function of AP coordinate *x* close to the midline (*y* = 0.2*R*). (*e*) Rate of strain in the DV as a function of the DV distance to ventral midline, along a transverse cut midway along AP (*x* = 0).

Using an energetic formulation, it can then be shown that Eq (3) is equivalent to the constrained minimisation:

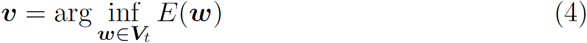
EQN4
where *E* is the rate of energy dissipation in the tissue, namely:

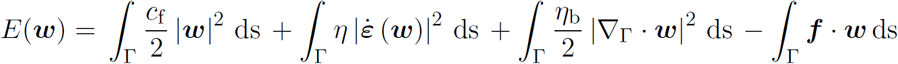
UEQN2

Our approach is to introduce a vector field ***θ*** that will act as a Lagrange multiplier to constrain the velocities ***v*** to be tangential. This field ***θ*** can be interpreted as the force needed to prevent normal deformations. In order to do this, we first define ***θ*** by ***θ*** = *γ**L***(***v***) where ***L***(***v***) = (***v*** · ***n***) ***y*** − *∇*_Γ_***v*** · ***n***, ***y*** is the curvature vector defined as ***y*** = Tr(*∇*_Γ_***v***)***n***, and *γ* is a strictly positive parameter. Then we note that ***V**_t_* = ker ***L***. The problem can now be rewritten as an unconstrained saddle-point problem:

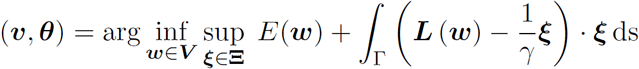
UEQN3
where *V* = (*H* ^1^ (Γ))^3^ and **Ξ** = (*L*^2^ (Γ))^3^, the set of square-integrable vector fields. We further introduce the surface pressure *p* which enforces the finite compressibility of the actomyosin in the tangential plane, *p* = −*η*_b_*∇*_Γ_ *· **v***. We can then write the problem as:

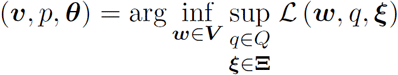
UEQN4
where *Q* = *L*^2^ (Γ),

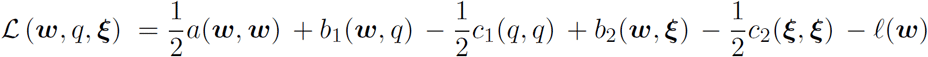
UEQN5
and *a, b*_1_*, b*_2_*, c*_1_ and *c*_2_ are the bilinear forms defined by:

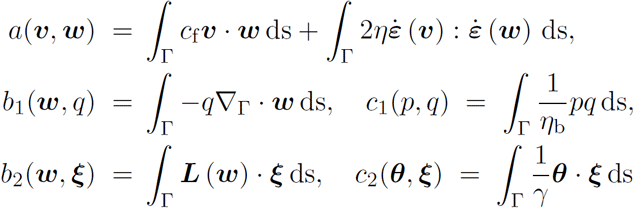
UEQN6
and 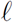 is the linear form define by 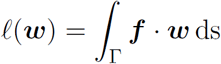. The saddle point can then be characterized as the solution of the linear problem:

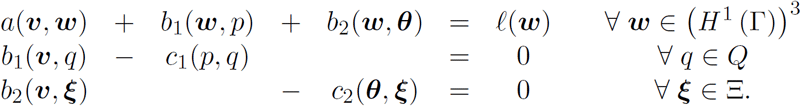
UEQN7

We show [25] that this problem has a unique solution in ***V*** × **Ξ** × *Q*.

## Mixed finite element approach

We solve the saddle point problem using the finite element method. This requires us to introduce a mesh Γ*h* approximating Γ. We use a triangular tessellation of second order, i.e. elements are curved triangles described by a quadratic transformation and whose largest dimension is smaller than the mesh size *h*. This ensures that the distance between any point of Γ*h* and Γ is at most *Ch*^3^, where *C* is a constant independent of *h*. Using this mesh, we define discrete functional spaces ***V**_h_*, **Ξ***_h_* for vector fields ***v**_h_*, ***θ**_h_* and *Q_h_* of scalar field *p_h_*. We approach the saddle point problem using the following formulation:

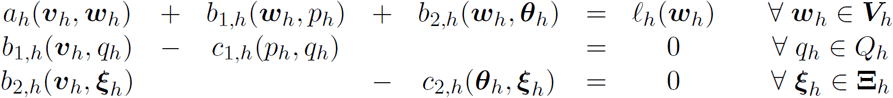
UEQN8
where *a_h_, b_1,h_, b_2,h_, c_1,h_* and *c_2,h_* are bilinear forms approximating the original forms, defined by:

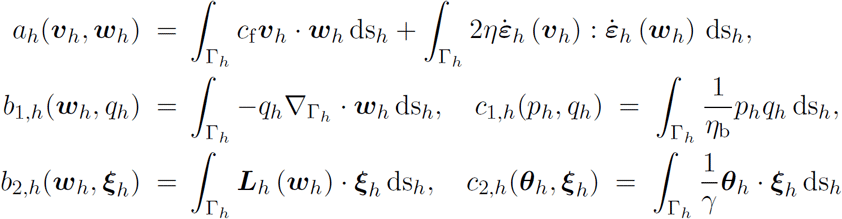
UEQN9
and 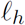 is the linear form defined by 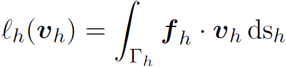. The choice of the finite element spaces cannot be made arbitrarily because it is a mixed problem.

It requires a suitable choice in order for the discrete problem converges towards the saddle point problem. Indeed, the discrete problem must verify two conditions called *inf–sup* or *Brezzi–Babuska* conditions (see [S3]): first between the spaces ***V**_h_* and *Q_h_* through the bilinear form *b*_1_*_,h,_* then between the spaces ***V**_h_* and **Ξ***_h_* through the bilinear form *b*_2_*_,h_*. In the absence of theoretical results on spaces that may verify these conditions, the idea is to produce compatible mixed finite element combinations in order to obtain the convergence. For this, we guided our choice by similarity with choices for which *inf–sup* conditions are verified in the case of classical problems (such as the three-dimensional Stokes problem).

## Numerical validation

Next, finite element spaces ***V**_h_, Q_h_* and **Ξ***_h_* must be specified. We base them on a triangular tesselation of the surface Γ (see next section) and choose Lagrange finite elements of degree 3 for ***V**_h_*, 2 for *Q_h_* and **Ξ***_h_*. We then check that this choice leads to a convergent approximation of the solution of Eq (3). In order to do so, we make an arbitrary choice of a velocity field on an arbitrary surface (a sphere), and calculate analytically the prestress that would be needed to achieve such a velocity field. We then run simulations on a series of meshes of decreasing triangle size *h* and monitor the evolution of the error ***v** − **v**_h_*. We show [25] that this decreases quadratically when *h* decreases, leading to pointwise errors (i.e., in *L^∞^* norm) smaller than 10*^−^*^3^ for all meshes of more than 10^4^ elements (*h* = 0.05). For these numerical tests, we chose: *c*_f_ = 10*^−^*^5^, *η* = 1, *η*_b_ = 10^3^ and *γ* = 10^7^.

**Fig S2:**
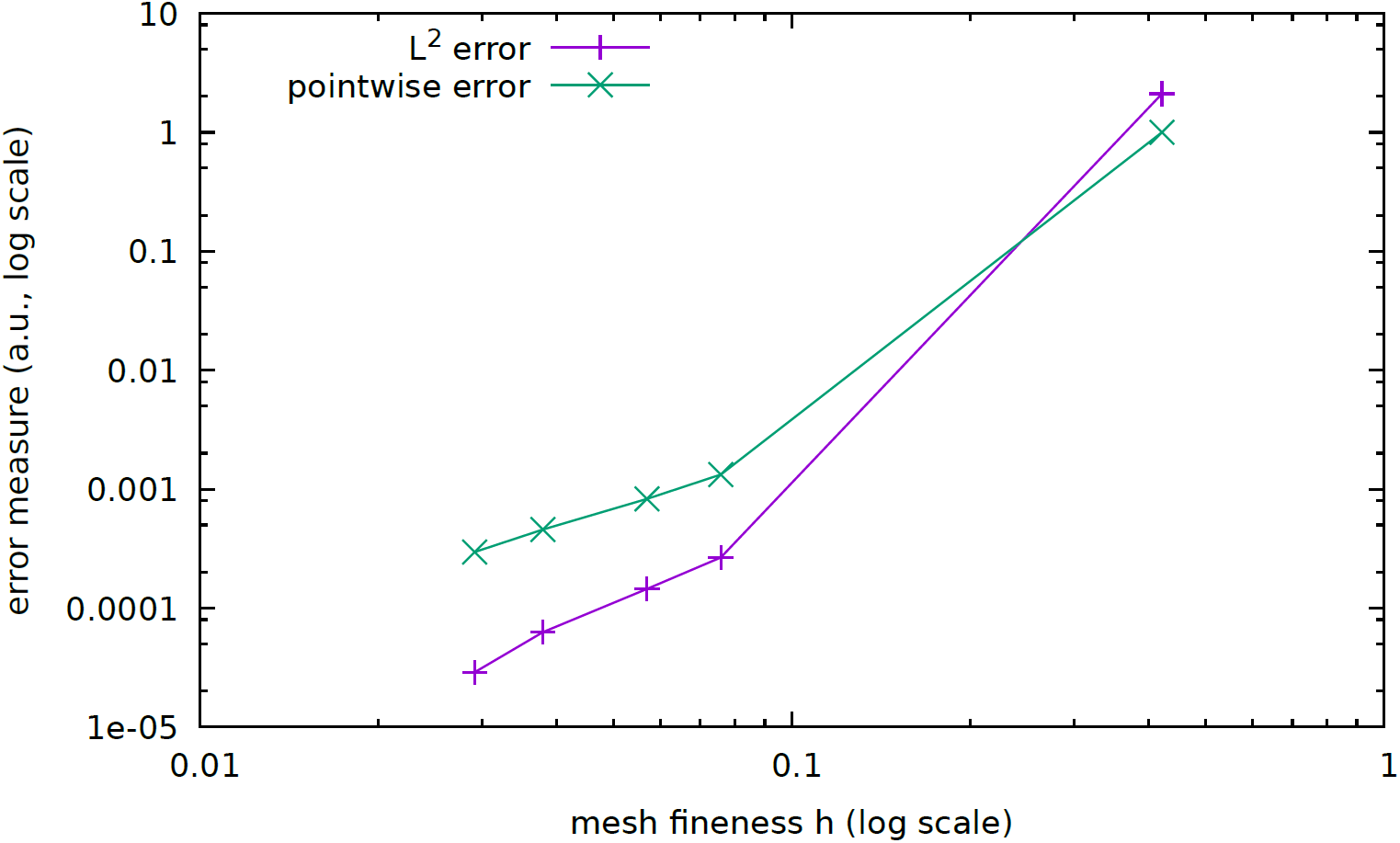
Convergence test. The mesh is refined (from right to left) and the error on an arbitrary flow field is seen to decrease. *L*^2^ error is the overall squared difference of calculated minus original velocity vectors, pointwise (*L_∞_*) error is the length of the largest difference between calculated and original velocity vectors over the whole mesh.

## Finite element mesh of the *Drosophila* embryo and resolution

We first describe the embryo shape with an analytical function, and then introduce a procedure to create a finite element mesh which will be fine enough to capture geometric details such as the cephalic furrow, while remaining of reasonable size in terms of the number of triangles (since the computational cost of our algorithms increases like *N* log(*N*) with number of triangles *N*).

The analytical function describing the embryo shape Γ = {*φ*(*x, y, z*) = 0} is chosen as:

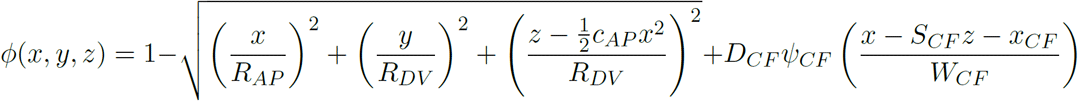
UEQN10
where *R_AP_* is the half-length of the embryo in AP, *R_DV_* its maximum radius in a transverse cut, *c_AP_* a curvature parameter corresponding to the curvature of the main axis of the embryo (defined as the locus of the centre of all transverse cuts), and parameters indexed with *CF* correspond to the cephalic furrow.

When *D_CF_* = 0, the cephalic furrow is absent, and the geometry corresponds to an ellipsoid of major axis along *x*, with radius *R_AP_*, and minor axes along *y* and *z* of equal radii *R_DV_*. The curvature parameter flattens the dorsal side (*z >* 0).

We take *R_DV_* = 1 as the reference adimensional length, *R_AP_* = 3*R_DV_* and *c_AP_* = 0.1*/R_DV_*, which leads to a shape close to the one of actual embryos.

The cephalic furrow depth is described by *D_CF_* = 0.1*R_DV_*, its position along the *x* axis in the mid-coronal plane *z* = 0 is given by *x_CF_* = −1.2*R_DV_*, and its inclination with respect to the (*y, z*) transverse planes is set by *S_CF_* = 0.3. The cephalic furrow has a total width *W_CF_* = 0.1*R_DV_* (exaggerated compared to real embryos, since a very thin and sharp feature would increase tremendously the computational cost), its shape is described by the function

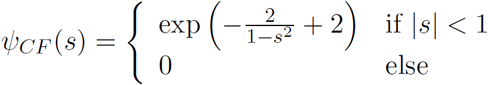
UEQN11
which is infinitely derivable, leading to a very smooth profile.

The mesh generation is delegated to mmgs software [S4], and the meshes used have around 46000 elements. The numerical resolution of the problem on this mesh is implemented in the open-source free software environment rheolef [S5].

## Microscopy and cell tracking

In Fig 1*a* we present data extracted from the tracking of GB extension in a wildtype embryo with the whole-membrane markers *resille-GFP* and *spider-GFP*, as described in [6]. Imaging was done using multidirectional selective plane illumination microscopy (mSPIM) [26], embryos were rotated to image four perpendicular views which were reconstructed into a whole embryo image stack post acquisition [S6]. Image stacks were acquired every 30 seconds.

In order to monitor the displacement of each cell, we first sensed the shape of the surface of the embryo in the three-dimensional mSPIM *z*-stack. We applied a grey-scale threshold to binarise the *z*-stack, highlighting only cell membrane-labelled signal. A ‘blanket’ of a fine meshwork of line-segments was dropped in the positive *z* direction down onto the embryo, until caught by cell membrane signal. This described the surface of the embryo accurately with a curved mesh, located at the apices of the embryonic epithelium. We used this embryo surface ‘blanket’ to extract curved image layers, with deeper layers shrinking progressively in single pixel steps in the direction normal to the local embryonic surface, towards the centre of the embryo [S7]. The radial depth giving the clearest view of the cell outlines of the embryonic epithelium was selected for tracking. Automated cell tracking with manual correction was performed using custom software written in ‘C’ and Interactive Data Language (IDL, Harris Geospatial). This tracked all cells in the chosen curved layer over time, identifying cell outline shapes and links forwards and backwards in time in an iterative process using an adaptive watershedding algorithm [14,15]. From each cell shape we calculated the location of the cell centroid (centre of mass). The coordinates of cell centroids, perimeter shapes, and links forwards and backwards in time were stored for all cells at each time point.

Using the relative movements of cell centroids, local tissue 2D rate-of-strain tensor (deformation rate) was calculated for small spatio-temporal domains focused on every tracked cell at each time point [15]. Local domains were composed of a focal cell and one corona of neighbouring cells over a 30 s interval (between successive frames). Local domains were first un-tilted according to the orientation of the local embryo surface and stretched to minimize artifacts caused by the local Gaussian curvature. All strain rates were then projected onto the embryonic axes, AP and DV. The rate of local area change was calculated as the trace of the 2D tissue strain rate tensor.

